# Computational Prediction of Heteromeric Protein Complex Disassembly Order with Hybrid Monte Carlo/Molecular Dynamics Simulation

**DOI:** 10.1101/2022.01.12.476000

**Authors:** Ikuo Kurisaki, Shigenori Tanaka

## Abstract

The physicochemical entity of biological phenomenon in the cell is a network of biochemical reactions and the activity of such a network is regulated by multimeric protein complexes. Mass spectroscopy (MS) experiments and multimeric protein docking simulations based on structural bioinformatics techniques have revealed the molecular-level stoichiometry and static configuration of subcomplexes in their bound forms, then revealing the subcomplex populations and formation orders. Meanwhile, these methodologies are not designed to straightforwardly examine temporal dynamics of multimeric protein assembly and disassembly, essential physicochemical properties to understand functional expression mechanisms of proteins in the biological environment. To address the problem, we had developed an atomistic simulation in the framework of the hybrid Monte Carlo/Molecular Dynamics (hMC/MD) method and succeeded in observing disassembly of homomeric pentamer of the serum amyloid P component protein in experimentally consistent order. In this study, we improved the hMC/MD method to examine disassembly processes of the tryptophan synthase tetramer, a paradigmatic heteromeric protein complex in MS studies. We employed the likelihood-based selection scheme to determine a dissociation-prone subunit pair at each hMC/MD simulation cycle and achieved highly reliable predictions of the disassembly orders with the success rate over 0.9 without *a priori* knowledge of the MS experiments and structural bioinformatics simulations. We similarly succeeded in reliable predictions for the other three tetrameric protein complexes. These achievements indicate the potential availability of our hMC/MD approach as the general purpose methodology to obtain microscopic and physicochemical insights into multimeric protein complex formation.

## Introduction

Physicochemical entity of the biological phenomenon in the cell is a network of biochemical reactions. Such a reaction network is composed of multimeric protein complexes, and the biological activities are dynamically regulated by their assembly and disassembly. Then, physicochemical characterizations of their assembly and disassembly orders, and those of the molecular dynamics in the cell, have drawn much attention to obtain comprehensive understanding of biological phenomenon from the molecular level.

Mass spectroscopy (MS) experiments and multimeric protein docking simulations based on structural bioinformatics techniques are the two major methodologies, which can give molecular insights into assembly and disassembly processes of multimeric protein complexes^1–6^. These two proteomics approaches play complemental roles each other. However, as briefly reviewed below, we find that these available methodologies have technical restrictions so far and room to employ an additional complemental methodology if we aim to physicochemically understand the assembly and disassembly processes.

MS approaches have been extensively employed to identify the subcomplex populations emerging in assembly and disassembly processes of multimeric protein complexes.^1,4–6^ The macroscopic information such as stoichiometric relationships among subunits can give straightforward clues to deduce assembly and disassembly orders of multimeric protein complexes.

Meanwhile, the spatial resolution of MS observations is bound to the macromolecular level, thus being unsatisfactory upon examining the microscopic factors regulating assembly and disassembly processes, that is, atomistic interactions such as hydrogen bonds. Furthermore, it is a possible concern that MS experiments cannot observe thermodynamically unstable subcomplexes under the influence of specific MS experimental conditions. They should appear in the assembly and disassembly processes but could not be retained during MS observation periods.^5^

These technical limitations have been compensated by the structural bioinformatics simulations to some extent. They can examine the assembly and disassembly orders based on atomic structures of proteins, and could theoretically give a complete sequence of subcomplex populations emerging through the assembly and disassembly processes.^2,3^ Assigning MS observed subcomplexes with such theoretical prediction can give global descriptions of assembly and disassembly processes and, furthermore, microscopic insights into the intermolecular interactions. Besides, in the structural omics era of today^7^, the data intensive bioinformatics studies are the promising approach to comprehensively explore potential protein assemblies, which should facilitate MS experimental researches.

Nonetheless, the complementary usage of these two proteomic approaches appears to be still unsatisfactory if we aim to physicochemically elucidate the formation mechanisms of multimeric protein complexes. Theoretical prediction by structural bioinformatics simulations is restricted to discuss static configurations of subcomplexes. Elucidation of their temporal dynamics during assembly and disassembly processes is essential to obtain deeper insights into the formation mechanisms. However, the structural bioinformatics simulations cannot be directly employed for subsequent physicochemical analyses such as free energy profile calculations.

Subcompelx formations in the cell should progress under the influence of both unbound subunits and molecules consisting of crowding milieu such as ions and metabolites^8,9^. Meanwhile, the structural bioinformatics simulations are designed to only give atomic coordinates of subcomplexes with ignoring those of these environmental molecules. Therefore, besides the structural bioinformatics approaches, we need another complemental approach to satisfactorily provide physicochemical and microscopic insights into formation mechanisms of multimeric protein complexes.

Under these circumstances, we recently proposed a hybrid configuration bias Monte Carlo/Molecular Dynamics (hcbMC/MD) approach and applied it to simulating the disassembly process of serum amyloid P component (SAP) homo pentamer.^10^ The atomistic simulation results we obtained were consistent with the observations of the earlier MS and structural bioinformatics study^6^ with regard to the emergence order of subcomplexes in the pentamer disassembly processes.

Furthermore, we discovered a novel preliminary dissociation event, the ring-opening reaction of the pentamer, and also analyzed the free energy profiles of the ring-opening reaction to show the possibility to experimentally observe the microscopic event. These achievements encourage us to suppose that our hcbMC/MD approach provides a complementary methodology for the MS and structural bioinformatics ones.

In this study, we extend application of the hcbMC/MD approach to heteromeric protein complexes, with the aim of developing it toward a general purpose methodology to examine disassembly processes of multimeric protein complexes. We begin with the consideration of tryptophan synthase tetramer as a representative heteromeric complex due to the following two reasons. Firstly, this tetramer has been extensively studied by MS approaches^6,11^ and knowledge of the disassembly order is available to test the performance of our hcbMC/MD simulations. Secondly, it should be more challenging to predict disassembly order of heteromeric complex than that of homomeric ones such as SAP pentamer (the reason will appear later). Thus, if we succeed in highly reliable prediction of the disassembly processes of heteromeric complexes, we can show methodological generality of our hcbMC/MD approaches.

Notably, the intrinsic challenge which we confront in this study is attributed to the heterogeneity of interfaces formed in the multimeric protein complex. In the previous study on SAP homo pentamer, each of the five interfaces is structurally equivalent, so that the order to dissociate a subunit pair in hcbMC/MD simulations does not affect generation of subcomplex populations (**Figure S1**). Meanwhile, the tryptophan synthase tetramer consists of two α subunits and two β subunits (**Figure 1**). The three interfaces in the tryptophan synthase tetramer are classified with the two groups: the two and the remaining are composed of the αβ and ββ subunit pairs, respectively.

**Figure 1.**
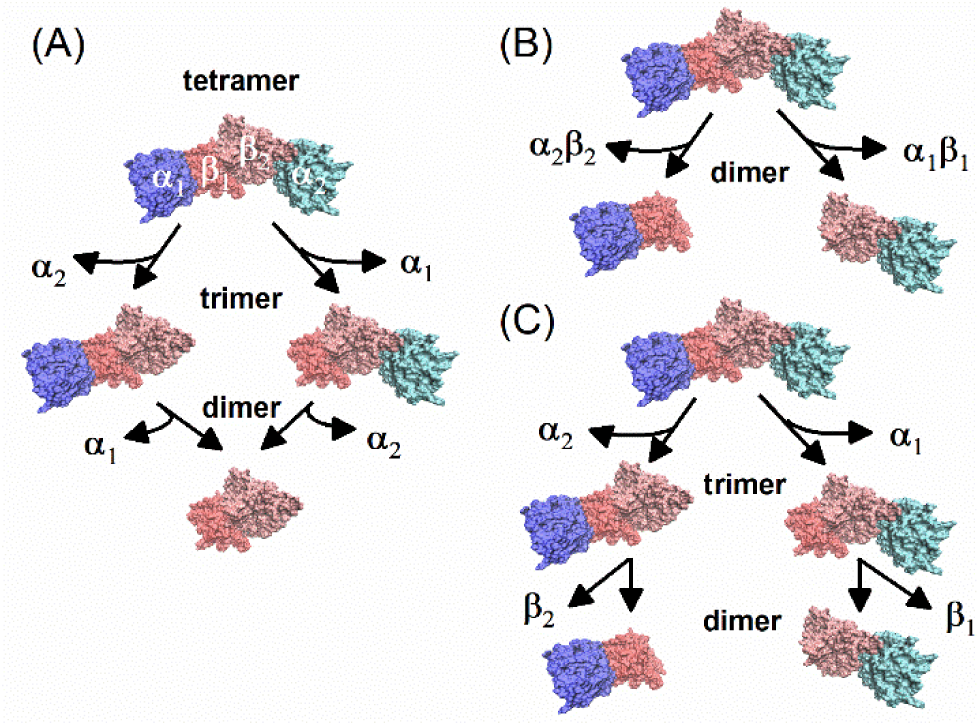
Disassembly pathway of tryptophan synthase tetramer. (A) ββ dimer formation. (B) direct αβ dimer formation. (C) αβ dimer formation via αββ/ββα trimer. Each of tryptophan synthase subunits is annotated by greek alphabet with a digit.

According to the mass spectroscopy (MS) studies, this tetramer disassembles into ββ dimer via subsequent dissociations of the two α subunits (**Figure 1A**). Thus, to reliably predict the disassembly order of tryptophan synthase tetramer with the hcbMC/MD approach, we need to dissociate a subunit pair via the proper order. However, the previous version of hcbMC/MD approach employs random selection of reaction coordinates at each cycle. Then, it is concerned that an unexpected subcomplex such as the αβ dimer (**Figure 1B** and **1C**) could appear with a certain frequency and substantially diminish the success rate to generate experimentally observed subcomplex populations (**Figure 1A**).

Recalling the possible problem upon applying the original hcbMC/MD approach to the tryptophan synthase tetramer, we propose a new reaction selection scheme from the point of methodological improvements. We focus on such an MS observation for stability of protein-protein interactions that reducing the number of salt bridge (SB) formation leads to the lowering of activation barrier of inter-subunit dissociation.^6^ To preferentially choose a dissociation-prone subunit pair, at each hcbMC/MD cycle without any *a priori* knowledge of MS and structural bioinformatics studies, we newly propose the likelihood-based selection scheme, with which a dimer of subunits with the smaller number of SB formation is more likely selected as a subunit pair to dissociate.

We test both this likelihood-based selection scheme and the previous one, in which a reaction pair is randomly selected at each hcbMC/MD cycle, and examine the simulation performance to predict disassembly order in a manner consistent with MS and structural bioinformatics approaches. Besides, hcbMC/MD simulations are employed for additional three heteromeric tetramers to assess the generality of our approach.

## Materials and Methods

### Setup of heteromeic tetramer systems

A set of atomic coordinates of each heteromeric tetramer^12–15^ was obtained from Protein Data Bank^16^ and the molecular components of the systems are summarized in **Table 1** (additionally, the amino acid sequences of the subunits are given in **Table S1**). All ligand molecules except for crystal waters were removed from the set of atomic coordinates. The atomic coordinates unresolved experimentally were complemented by using Swiss Model.^17,18^ Each of the histidine residues was in the Nε protonation state, and all of the aspartate and glutamate residues has the carboxyl group in the deprotonated state, where we consider physiological pH value in the cell. Each tetramer structure was solvated in the rectangular box and was electrically neutralized by adding counter ions. Besides, we prepared the SAP pentamer system in the manner similar to our previous study.^10^ The simulation details are given in **SI-1** in Supporting Information.

**Table 1.** Molecular components of tetramer systems.

| proteins | number of amino acid<br>residue | number of cosolute and<br>solvent molecules |  | PDB ID |
| --- | --- | --- | --- | --- |
|  |  | K<sup>+</sup> | water |  |
| Tryptophan Synthase | 1326<br>(268/395 for $\alpha/\beta$ subunit) | 30 | 93139 | 4HPX |
| Acetyl-coenzyme A carboxylase<br>carboxyl transferase | 1138<br>(314/255 for $\alpha/\beta$ subunit) | 22 | 76295 | 2F9I |
| Co-type nitrile hydratase | 838<br>(200/219 for $\alpha/\beta$ subunit) | 28 | 88303 | 3QXE |
| Anthranilate synthase | 1418<br>(517/192 for $\alpha/\beta$ subunit) | 32 | 88268 | 1I7Q |

To calculate the forces acting among atoms, AMBER force field 14SB^19^, TIP3P water model^20,21^, and Joung-Cheatham ion parameters adjusted for the TIP3P water model^22,23^ were employed for amino acid residues, water molecules, and ions, respectively. Molecular modeling of each system was performed using the LEaP modules in AmberTools 17 package^24^. Molecular mechanics (MM) and molecular dynamics (MD) simulations were performed under the periodic boundary condition with GPU-version PMEMD module in AMBER 17 package^24^ based on SPFP algorithm^25^ with NVIDIA GeForce GTX1080 Ti.

We performed the 10-ns NPT MD simulation for each system and employed the snapshot structure at 10 ns for the initial atomic coordinates in the following hcbMC/MD simulations. We confirmed that each of these initial structures shows characteristic inter-subunit contacts under thermal fluctuation (see **SI-2** in Supporting Information, and compare **Tables 2** and **Table S2**). The simulation procedure is similar, except for the molecular system, to that of our previous study^10^ so that we give the related information in **SI-3** and **SI-4** in Supporting Information.

**Table 2.** Inter-subunit salt bridge formation in the initial atomic coordinates for the hybrid configurational bias MC/MD simulations.

| tetramer | subunit pair |  |  |  |  |  |
| --- | --- | --- | --- | --- | --- | --- |
| | $\alpha_1\beta_1$ | $\alpha_2\beta_2$ | $\alpha_1\alpha_2$ | $\beta_1\beta_2$ | $\alpha_1\beta_2$ | $\alpha_2\beta_1$ |
| Tryptophan Synthase | 6 | 6 | 0 | 9 | 0 | 0 |
| Acetyl-coenzyme A carboxylase<br>carboxyl transferase | 15 | 15 | 0 | 0 | 12 | 13 |
| Anthranilate synthase | 21 | 28 | 12 | 0 | 0 | 0 |
| Co-type nitrile hydratase | 30 | 33 | 0 | 3 | 0 | 0 |

### Hybrid configuration bias MC/MD simulation scheme

The original and newly proposed hcbMC/MD schemes are illustrated in **Figure 2A** and **2B**, respectively. We here explain the workflow of hcbMC/MD simulations and give further details in the next subsection. The simulation procedures of configuration generations are similar to those in our previous study^10^, so that they are given in **SI-5** in Supporting Information. Each annotation assigned to computational operation (*e.g.*, **1. Relaxation stage**) is common between the illustration in **Figure 2** and the following descriptions.

**Figure 2.**
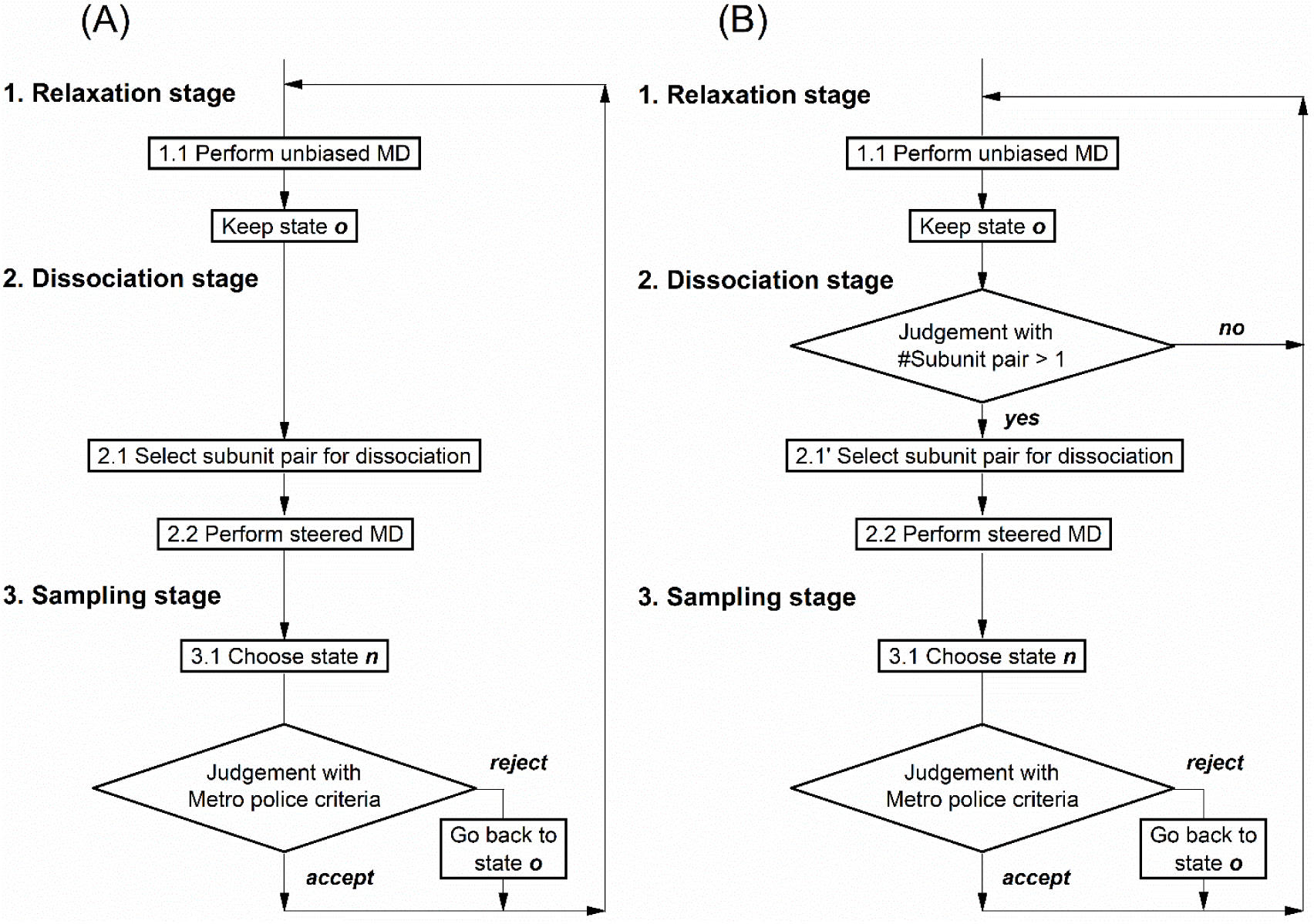
Workflows of configuration bias hybrid Monte Carlo/Molecular Dynamics simulations. (A) Original scheme proposed in our previous study^10^. (B) The new scheme with the likelihood-based selection of reaction coordinate.

#### 1. Relaxation stage

**1.1** Generate *M* trial configurations {**a**_1_,**a**_2_,…**a***_M_*} by an unbiased MD simulation.

We assign the last snapshot obtained from the simulation ( **a***_M_* ) as an old configuration, denoted by **x***_o_* (state ***o***).

#### 2. Dissociation stage

**2.1** *The previous hcbMC/MD scheme*: randomly choose a pair of subunits in the *_N_*C_2_, combination of the two among the *N* subunits (where the value of *N* is set to 4, due to considering the tetramers in this study), candidates at each hcbMC/MD cycle.

**2.1’** *The newly proposed hcbMC/MD scheme*: search subunit pairs which form inter-subunit salt bridges and pick up one among them by using the likelihood function (the formula appears in the next subsection).

If we have only one candidate via the search process at an hcbMC/MD cycle, move to **1.1** and start a new unbiased MD simulation by using **x***_o_* (see **Figure 2B**).

**2.2** Generate *M* trial configurations {**b**_1_,**b**_2_,…**b***_M_*} by a steered MD simulation starting from the **x***_o_* .

The pair of subunits is dissociated by imposing an external force on inter-center of mass distance. A target value of the distance was set to *d*_0_ + Δ*d*, where *d*_0_ and Δ*d* are an initial value of the distance calculated from the **x***_o_* at the cycle and an incremental distance (taken as an integer), respectively.

#### 3. Metropolis trial

**3.1** Calculate the Rosenbluth factors for the trial configurations, {**a**_1_,**a**_2_,…**a***_M_*} and {**b**_1_,**b**_2_,…**b***_M_*} , referred to as *W*(*o*) and *W*(*n*) hereafter, respectively:

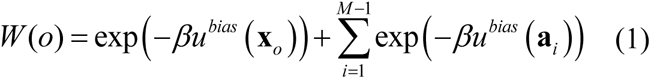

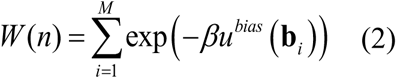

The bias energy function *u^bias^* will be specified later. *β* denotes inverse of k_B_T, where k_B_ and T are Boltzmann constant and system temperature, respectively. We select one configuration among {**b**_1_,**b**_2_ ,…**b***_M_* } , denoted by **x***_n_* (state ***n***), with a probability

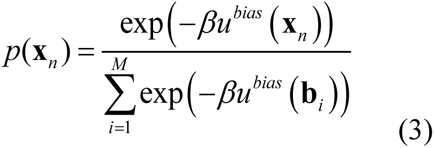

**3.1** The configuration change is accepted with a probability

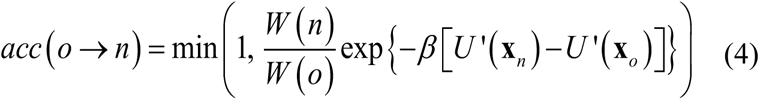

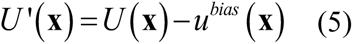

where **x** and *U* (**x**) denote a configuration and the native potential energy function of the system with configuration of **x** , respectively.

If the **x***_n_* is accepted, the next cycle starts from this configuration. If not, **x***_o_* is used for the next cycle.

#### Weighted function for configuration selections and likelihood function for reaction pair selections

As in the case of our previous study,^10^ the bias energy to calculate the Rosenbluth factor, *u^bias^*, is a function of the total number of inter-subunit native contact formed in a multimeric protein complex (*n_NC_*), where the initial atomic coordinates of the hcbMC/MD simulations are used as the reference structure to define the native contacts:

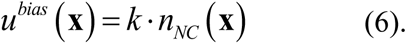

**x** denotes a set of atomic coordinates of the system. The coefficient *k* is set to *β*^−1^ at 300 K, that is 0.6 [kcal/mol] by assuming energetic stabilization obtained from hydrophobic atomic contacts^26,27^.

In the hcbMC/MD simulations with the newly proposed likelihood-based scheme, the subunit pair undergoing (partial) dissociation at each cycle is selected among the pairs which have nonzero number of inter-subunit salt bridge formation (the definition is given later). A pair of subunits *S_i_* and *S_j_*, or *S_i_S_j_*, is weighted by the likelihood function, *p* (*S_i_S_j_*|{*N_SB_* }):

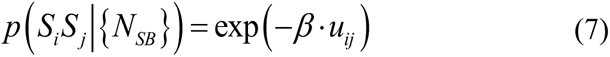

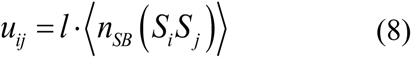

where *n_SB_*(*S_i_S_j_*) and the bracket respectively denote the number of salt bridge formed between subunits S*_i_* and S*_j_* in a snapshot structure and averaging operation over all *M* snapshots obtained from an unbiased MD trajectory at the given cycle. {*N_SB_*} is the set of ⟨n*_SB_* (*S_i_S_j_*)⟩ calculated for each of the pair of subunits. Thus, *p* (*S_i_S_j_*|{*N_SB_* }) is defined as such a relative probability that a pair of subunits *S_i_* and *S_j_* is selected under {*N_SB_*} . The coefficient *l* in eqn. 3 is set to 1 kcal/mol as test, and *β* is the inverse temperature at 300 K. Of note, we designed this likelihood function to preferentially select a subunit pair with smaller (but nonzero) number of salt bridge formation, which most likely dissociates at an hcbMC/MD cycle.

### Trajectory Analyses and Bayesian Inference

Atomic contacts and salt bridge formations were analyzed with the cpptraj module in AmberTools 17 package^24^. As in our previous study^10^, atomic contacts were calculated by using the non-hydrogen atoms and the native contacts are defined by considering the initial atomic coordinates for the hcbMC/MD simulations. The distance criterion for atomic contacts was set to 3.5 Å. We find a subunit pair configurationally dissociated when there are no atomic contacts between the subunit pair.

Besides, we defined the following quantities by using the number of atomic contacts between a subunit pair:

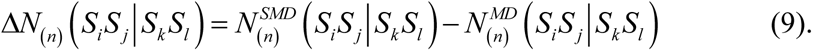

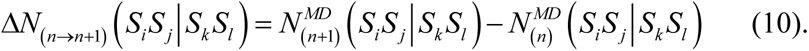

*n* denotes the hcbMC/MD cycle number and we here assume that Metropolis trial is accepted at the *n*^th^ cycle. *S_i_*, *S_j_*, *S_k_* and *S_l_* are assigned to any of the four subunits: α_1_, α_2_, β_1_ and β_2_. A digit in lower case is given to a subunit annotation, when two same type of subunits are individually discussed. It is noted that *S_i_S_j_* and *S_k_S_l_* in the parentheses are separated by the bar. A subunit pair on the left side of the bar (that is, *S_i_S_j_* in eqn. 9 and eqn. 10) is for calculation of the number of atomic contacts, while that on the right side of the bar (that is, *S_k_S_l_* in eqn. 9 and eqn. 10) is for dissociation reaction pair in the SMD simulation at the *n*^th^ cycle.

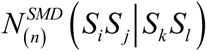 means the number of atomic contact calculated from the Metropolis-accepted SMD snapshot structure, while 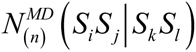 means the number of atomic contact calculated from a snapshot structure obtained from the unbiased MD simulation at the *n*^th^ cycle.

In particular, if *S_i_S_j_* is equal to *S_k_S_l_*, we used shortened representations:

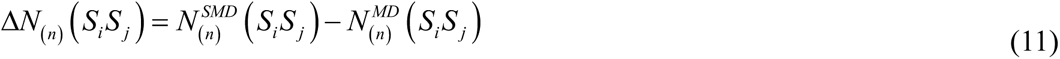

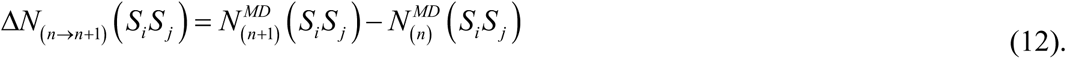

The salt bridge (SB) formation was defined at the atom-pair level, with regard to the side chains of Arg, Lys, Asp, and Glu residues. We suppose the SB formation if an atomic distance between X and Y is smaller than 3.5 Å, where X and Y denote a nitrogen atom bound to hydrogen atom in either Lys or Arg residue and an oxygen atom in either Asp or Glu residue, respectively. In the present case, we employed this SB definition to distinguishably characterize geometrical differences among subunit interfaces and relevantly select dissociation-prone subunit pair with eqn. 7.

We used Bayesian inference to estimate 95% credible interval of the success rate of expected subcomplex emergence, *θ*. This binary process is modeled as follows: The prior distribution is the uniform distribution and the likelihood is given as a binomial distribution, *_N_C_k_* ·*θ*^k^·(1−*θ*)*^N^*^-k^, where *N* and *k* denote the total number of hcbMC/MD simulation trials (that is 20 in this study) and the number of trials giving an expected subcomplex. Markov chain Monte Carlo simulations to estimate the rate are repeated 200,000 times by using independent two chains. The first 5000 data are discarded and then samples are collected by 10 intervals. Finally, 38000 sampled data were used for estimation of *θ* in total. We selected this procedure according to the reference of Bayesian inference.^28^

Bayesian inference was carried out with WinBUGS^29^ plus R^30^. Molecular structures were illustrated using Visual Molecular Dynamics (VMD).^31^

## Results and Discussion

### The hybrid MC/MD approach can reliably predict disassembly processes of heteromeric protein complexes

Firstly, we performed the independent 20 hcbMC/MD simulations with the ‘likelihood-based selection (LS)’ scheme to simulate disassembly processes of tryptophan synthase tetramer (TS). An increment of inter-center of mass (COM) distance for SMD simulations, annotated by Δ*d*, is sampled from the integer distribution which ranges from 13 to 17 Å at each hcbMC/MD cycle as test. Each of hcbMC/MD simulations was extended by 500 cycle (100 ns in total) at maximum.

As remarked above, the MS experiment shows that the tryptophan synthase tetramer is decomposed into two α subunits and ββ dimer complex, finally (**Figure 1A**). Meanwhile, it is possible that another subcomplex species, αβ dimer (**Figure1B** and **1C**), emerges via an hcbMC/MD simulation. Since the emergences of ββ dimer and αβ dimer are mutually exclusive, we can evaluate the performance of hcbMC/MD approach by focusing on the simulation cycle at which a dimer species first emerges, referred to as χ^dimer^, hereafter. Then, in the following analyses, we consider hcbMC/MD cycles by the χ^dimer^ for each of the simulations and calculate the success rate for emergence of ββ dimer via the simulations, referred to as *θ*_ββ_.

The simulation results are summarized in **Table 3**. The 20 hcbMC/MD simulations with the LS schemes give *θ*_ββ_ of 0.90 (the representative configuration of ββ dimer is shown in **Figure 3A** and subcomplex generations along the hcbMC/MD cycle are discussed in **Figure S2**). With this sample size, the Bayesian inference for θ_ββ_ shows that the success rate is credibly high compared with 0.5 (**Figure 4**). Then, it could be expected that our hcbMC/MD simulations give the MS-consistent disassembly order of tryptophan synthase tetramer with sufficiently high success rate.

**Figure 3.**
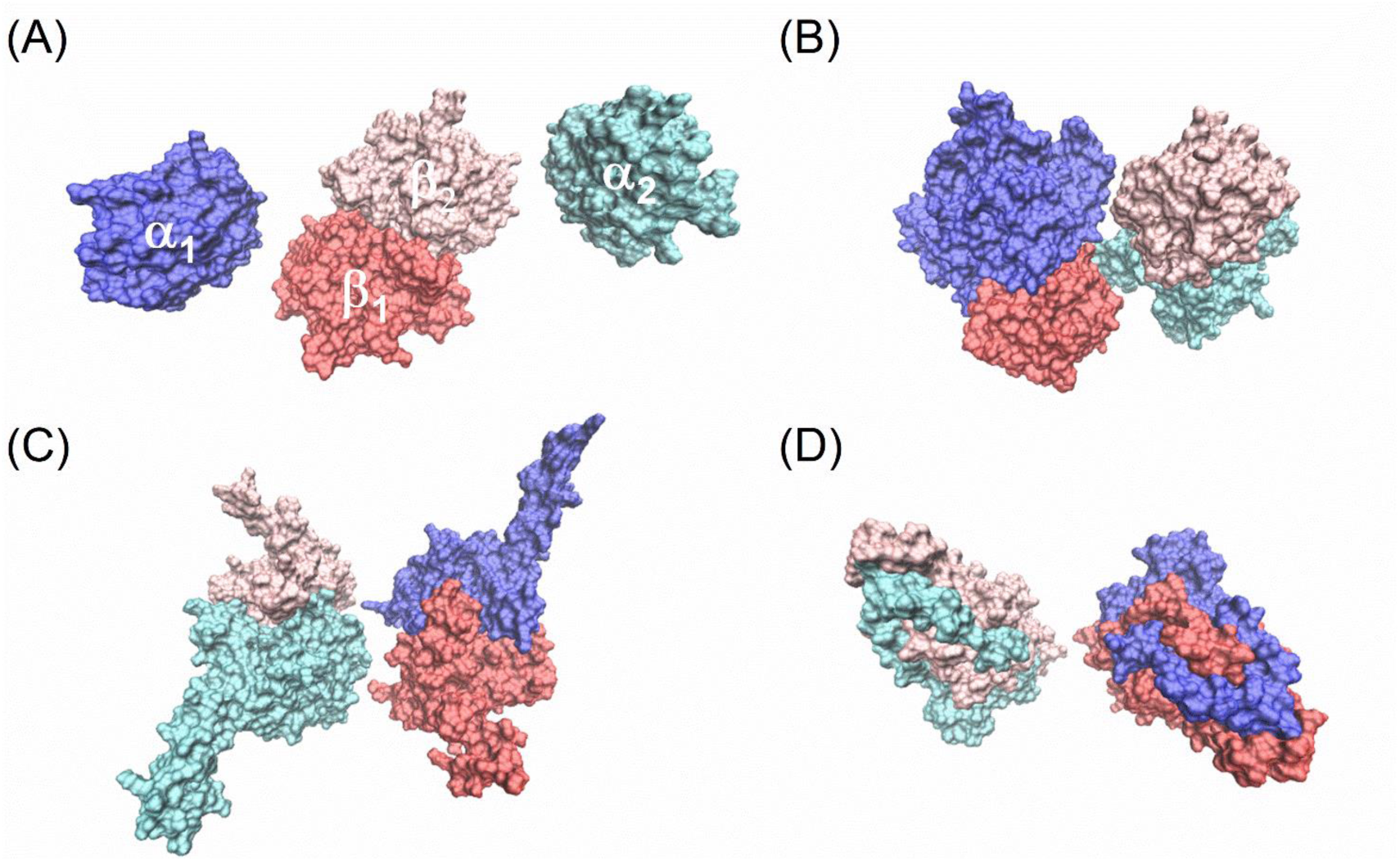
Representative structures of the expected dimer species, which are obtained by using the hcbMC/MD simulations. (A) Tryptophan synthase tetramer. (B) Acetyl-coenzyme A carboxylase carboxyltransferase. (C) Anthranilate synthase. (D) Co-type nitrile hydratase. In each panel, blue and cyan are for α_1_ and α_2_ subunits, while red and pink are for β_1_ and β_2_ subunits, respectively.

**Figure 4.**
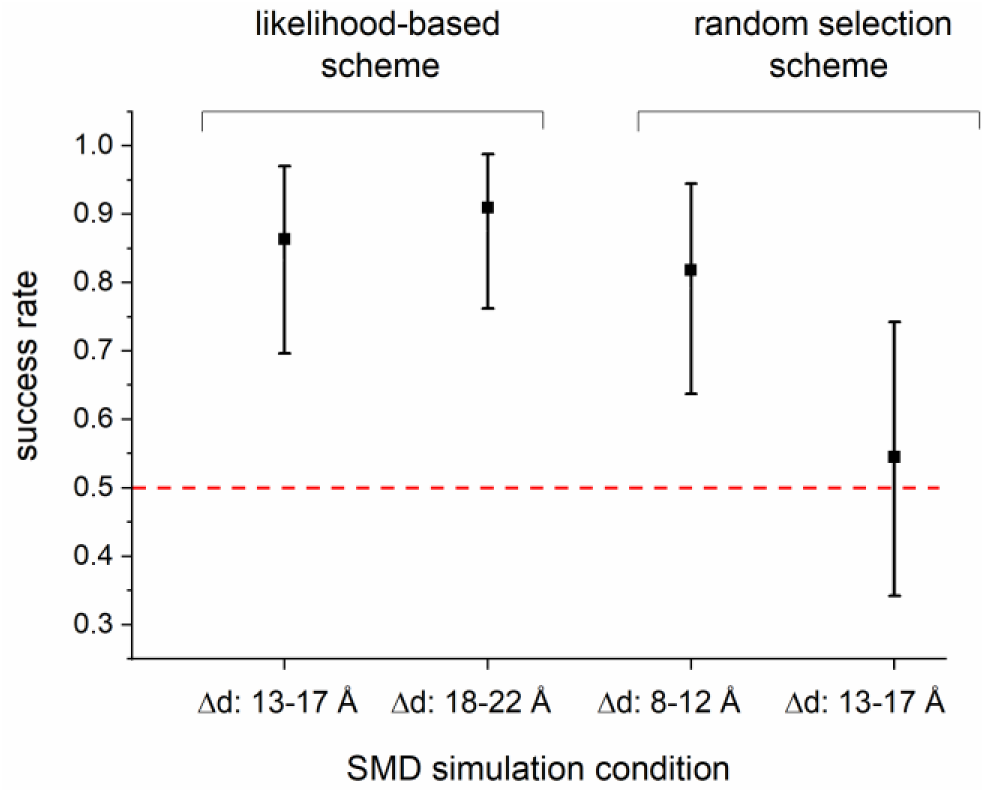
Bayesian inference of success rate of tryptophan ββ dimer emergence. Error bars denotes 95% credible interval and a square on the bar corresponds to the average value. Δ*d* means the range of increase of distance in a steered MD (SMD) simulation. The probability of 0.5 is highlighted by the red dotted line.

**Table 3.**
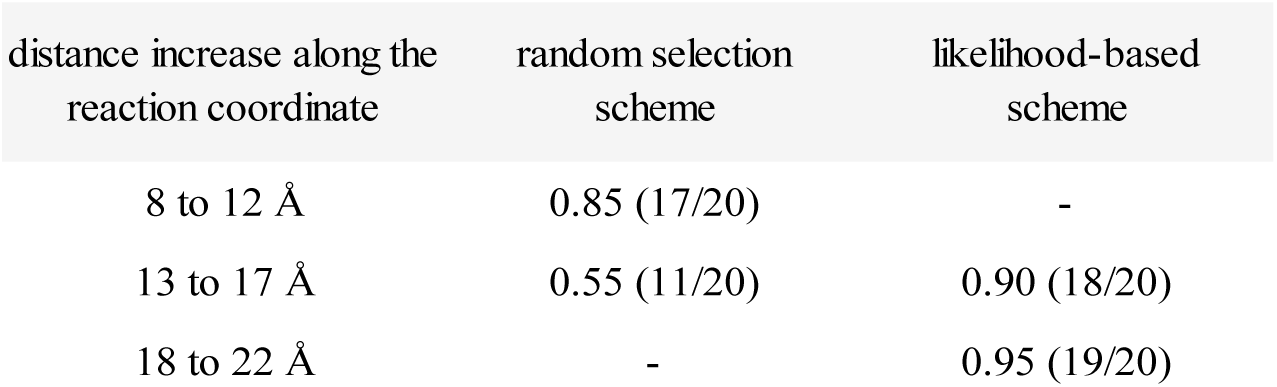
Success rate of ββ dimer emergence from the tryptophan synthase tetramer. The number of succeeded and total hcbMC/MD simulations are found at the left and right sides across the slash symbol in the parentheses.

We could suppose that such a high emergence rate of β_1_β_2_ dimer is due to use of the LS scheme. Actually, as shown below, we succeeded in preferably selecting the αβ dissociation in hcbMC/MD simulations (in other words, we preferably suppress selection of ββ dissociation.) By the first emergences of dimer species, χ^dimer^, the 583 SMD simulations in total were carried out within the 20 hcbMC/MD simulations. Either α_1_β_1_ or α_2_β_2_, and β_1_β_2_ are 531 and 52 times selected as the dissociation target in SMD simulations (91% and 9%), respectively. Meanwhile, the random selection (RS) scheme gives the selection frequencies similar among these subunit pairs (see **Table S3**, and the detailed discussions about hcbMC/MD simulations with the RS scheme will appear in the next subsection).

To quantitatively characterize availability of the LS scheme in suppression of selecting β_1_β_2_ dissociation, we examine a ratio of likelihoods for dissociating subunit pair selection

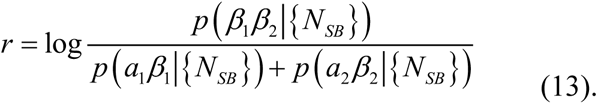

*p* (*S_i_S_j_*|{*N_SB_*}) in eqn. 13 is the likelihood function (while it is noted that *S_i_S_j_* is replaced with either of α_1_β_1_, α_2_β_2_ or β_1_β_2_ in the formula), whose definition is given in eqn. 7 (see Materials and Methods section). We can consider that a smaller value of *r* denotes lower probability of selecting β_1_β_2_ as a dissociation-prone subunit pair in an SMD simulation. It is the characteristic situation where *r* takes 0, the probability of selecting β_1_β_2_ dissociation is equal to that of selecting either α_1_β_1_ or α_2_β_2_ dissociation.

The 18 simulations, which successfully give β_1_β_2_ dimers, contribute to 472 of the 583 cases. In the 462 cases (97.9% of the 472), either α_1_β_1_ or α_2_β_2_ dimer is always selected. The values of *r* are lower than −2 in the 424 cases (92% of the 462). Under such a situation, the probability of selecting β_1_β_2_ dimer is 1% or smaller. It thus appears to be reasonable that the αβ dissociation is dominantly selected.

Meanwhile, β_1_β_2_ dissociation is only selected 10 times (2.1% of the 472) (**Figure 5A**). In the 10 cases of selecting β_1_β_2_ dimers, *r* takes −1 or greater (0.65 at maximum). Since this value range corresponds to 10% or greater in terms of probability of selecting β_1_β_2_ dimer, it is likely that β_1_β_2_ dissociation is chosen through probabilistic processes and employed for SMD simulations. Nonetheless, as for these simulations, each of dissociated snapshot structures is rejected at the Metropolis judgement, thus not working for β_1_β_2_ dissociation. In total, it can be said that selection of either α_1_β_1_ or α_2_β_2_ dissociation is sufficiently facilitated, then leading to generation of β_1_β_2_ dimer.

**Figure 5.**
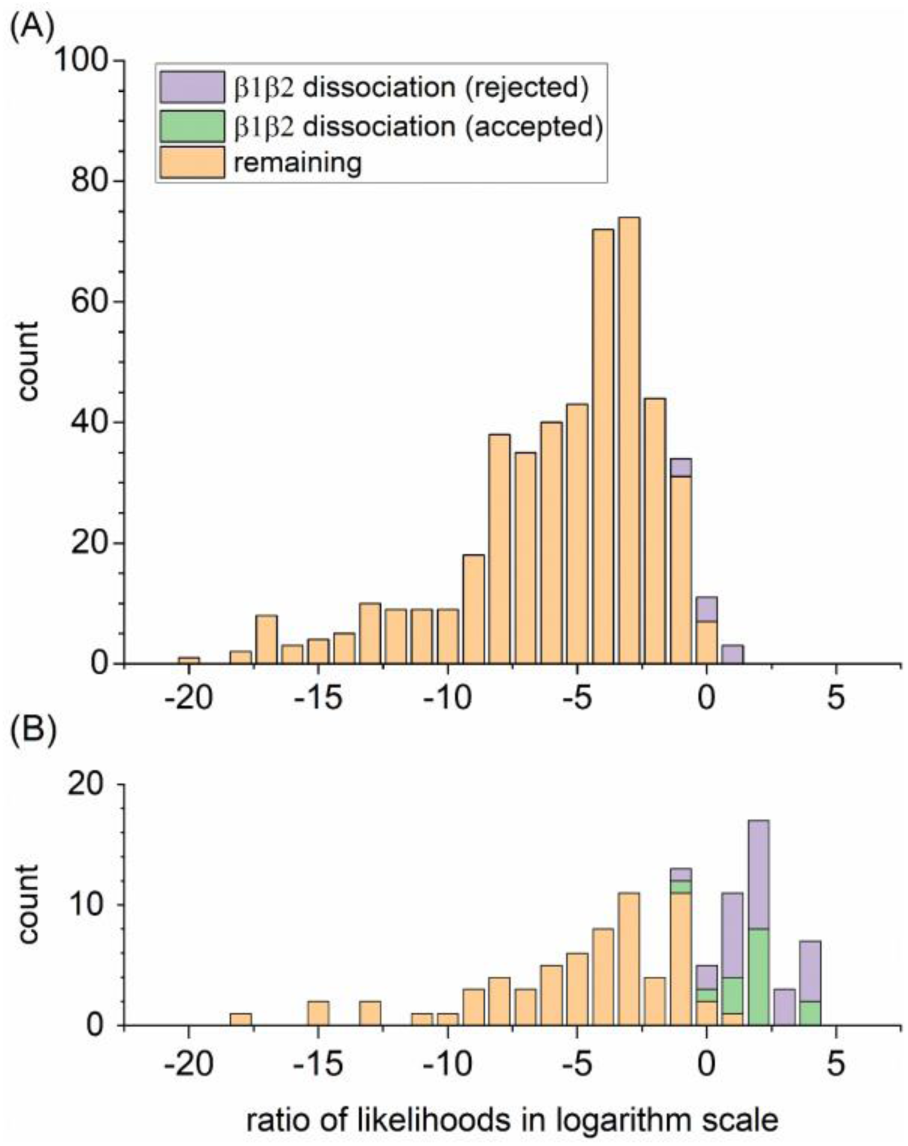
Distribution of relative likelihood of ββ dissociation to αβ dissociations. (A) The 18 simulations generating ββ dimer. (B) The 2 simulations generating αβ dimer. Orange bars are for either α_1_β_1_ or α_2_β_2_ dissociation selection. Green and purple bars are for Metropolis-accepted and -rejected ββ dissociation selection, respectively.

Meanwhile, there are the two failed hcbMC/MD simulations which give an unexpected αβ dimer, instead of the ββ dimer. These failed simulations undergo 111 SMD simulations in total when we consider hcbMC/MD cycles by χ^dimer^ for each of the two simulations. Selection of β_1_β_2_ dissociation is counted as 42 of 111 (37.5%), and 15 of the 42 are accepted via the Metropolis judgement (**Figure 5B**).

We can explain that such a relatively high frequency of selecting ββ dissociation, which is observed for the two simulations, is due to transient reduction of salt bridge formation between ββ under influence of thermal fluctuation (see the related discussion in **SI-6** in Supporting Information).

Nonetheless, we suppose that the effect of thermal fluctuation is satisfactorily suppressed in total. As remarked above, the success rate of 0.9 is credibly high to predict generation of the MS-observed ββ dimer, and emergence of the unexpected subcomplex appears to be relatively minor. This is probably due to preferential selection of αβ dissociation, which is supported by that we employ the average number of salt bridge formation (*N*_SB_). In the most cases, average values obtained under thermal fluctuation should relax the effect of transient change of *N*_SB_ at a single time point in the MD simulation.

It may be technically possible to further suppress the ββ selection by increasing the dependence of the likelihood function on *N*_SB_. Meanwhile, we suppose that such a modification is not necessarily required so far, because we already obtained a reliable prediction of subcomplex generation with the current formula of the likelihood function. Then we continue to use this likelihood function for the additional simulations discussed below.

We can furthermore show that this implementation of LS scheme straightforwardly improves the previous hcbMC/MD scheme. We similarly applied the newly proposed hcbMC/MD scheme to serum amyloid P component protein (SAP) and tested the reproducibility of our earlier study with regard to generation of the subcomplex population. We performed the 5 independent hcbMC/MD simulations for SAP system and found disassembly through each of two dissociation pathways, dimer-trimer formation and monomer-tetramer formation (see **Figure S3** and **SI-1** in Supporting Information). We also observed the preliminary event of the disassembly, SAP pentamer ring-opening reaction. Accordingly, we conclude that the usage of the LS scheme in hcbMC/MD simulations is a straightforward improvement of the original version of the hcbMC/MD scheme.

### Likelihood-based selection stably generates MS-predicted subcomplex population

Next, we performed the independent 20 hcbMC/MD simulations with the random selection (RS) scheme. As in the case of our previous study^10^, an increment of inter-COM distance at each hcbMC/MD cycle is sampled from the integer distribution which ranges from 8 to 12 Å. These simulations give 0.85 as the value of *θ*_ββ_ (**Table 3**). Similar to the above case, the Bayesian inference for *θ*_ββ_ shows that the success rate is credibly high compared with 0.5 (see **Figure 4**). We here observed generation of the ββ dimer in the 17 simulations. Among the three failed simulations, one has not reached a dimer generation, so that we excluded this simulation and discuss the 19 ones for the following analyses (subcomplex generations along the hcbMC/MD cycle are discussed in **Figure S4**).

Recalling the design of the RS scheme, this highly credible success rate for the ββ dimer generation is unexpected and rather opposite to an intuition from the technical viewpoint. In the RS scheme, we consider all possible pairs of subunits, that is _4_*C*_2_ = 6, to accelerate the dissociation reaction. At each hcbMC/MD cycle, any of the 6 pairs is randomly chosen with no regard to formation of protein-protein interface and structural characteristics of inter-subunit interaction. Actually, each of the six subunit pair is selected with the similar frequency over the 19 hcbMC/MD simulations (**Table S3**). It can be straightforward to think that this random selection for dissociating subunit pair should have given a lower success rate of expected ββ dimer generation, 0.5 or lesser (see **Figure S5**).

Then, we considered the mechanism for emergence of ββ dimer with the credibly high success rate, below. Recalling the structural observation that α_1_β_1_ and α_2_β_2_ form the smaller number of salt bridge between the subunits than β_1_β_2_ (**Table 2**),^6^ we can speculate that the number of atomic contacts between α and β subunits more rapidly decrease via SMD simulations than that between the two β subunits. We made the speculation by assuming that an inter-subunit interface with the greater number of atomic contacts are robust and easily recoverable even if the atomic contacts are partly broken by mechanical perturbation via SMD simulations.

We examine this speculation by analyzing Δ*N*_(_*_n_*_)_ (*S_i_S_j_*) and Δ*N*_(_*_n_*_→_*_n_*_+1)_ (*S_i_S_j_*) , which describe the quantitative change of the number of atomic contacts formed between subunits *S_i_* and *S_j_*. The former quantifies the effects of SMD simulations on break of atomic contact formation between subunits, while the later quantifies the total number of disrupted atomic contact formation after the recovery by the following unbiased MD simulation at the *n*+1^th^ cycle (see eqn 9 and eqn 10 in Materials and Method section for details of formula). It is noted that, at the *n*^th^ cycle annotated above, a snapshot structure obtained from the SMD simulation is accepted via Metropolis judgement and is employed for the initial atomic coordinates of the next *n*+1^th^ cycle.

Comparing Δ*N*_(_*_n_*_)_ (*S_i_S_j_*) among β_1_β_2_, α_1_β_1_ and α_2_β_2_, β_1_β_2_ shows the largest decrease in the number of atomic contacts (**Table 4**). This could be due to the larger size of β subunits, which have the greater number of atomic contacts on the interface area between β_1_ and β_2_, thus leading to the greater loss of atomic contacts by the SMD-driven dissociation. However, such decrease can be compensated by the following conformational relaxation. Actually, β_1_β_2_ shows the largest value of ΔN_(*n*→*n*+1)_ (*S_i_S_j_*) , the smallest disruption of atomic contacts by SMD simulations, which is 1.5-fold greater or more than those of α_1_β_1_ and α_2_β_2_. In total, we can suppose that atomic contacts formed between β_1_β_2_ are relatively robust compared with those formed between α_1_β_1_ and α_2_β_2_. The above observations indicate that dissociations of β_1_β_2_ relatively slowly progress compared with those of α_1_β_1_ and α_2_β_2_, then showing microscopic origin of retention of the dimer formation under the dissociating effect of SMD simulations.

**Table 4.** Breakage and recovery of atomic contact formation between subunit pair. Error estimation denotes 95% confidential interval.

| dissociated pair $S_i S_j$ | $\Delta N_{(n)}(S_i S_j)$ | $\Delta N_{(n \rightarrow n+1)}(S_i S_j)$ | # hcbMC/MD cycle for analyses |
| --- | --- | --- | --- |
| $\alpha_1\beta_1$ | $-12.16 \pm 0.02$ | $-4.39 \pm 0.02$ | 261 |
| $\alpha_2\beta_2$ | $-13.26 \pm 0.03$ | $-5.59 \pm 0.03$ | 225 |
| $\beta_1\beta_2$ | $-14.84 \pm 0.02$ | $-2.83 \pm 0.02$ | 277 |

Furthermore, dissociations of α_1_β_1_ and α_2_β_2_ can be accelerated by SMD simulations for a non-contacted subunit pair, α_1_α_2_, α_1_β_2_ or α_2_β_1_. As indicated by analyses of ΔN_(_*_n_*_→_*_n_*_+1)_ (*S_i_S_j_*|*S_k_S_l_*) in **Table 5**, α_1_β_1_ and α_2_β_2_ shows relatively large losses of atomic contact formation (2.26 and 2.47 at maximum, respectively) via SMD simulations for α_1_β_2_ and α_2_β_1_ dissociations, respectively. Meanwhile, β_1_β_2_ shows robustness and recovery of atomic contact formation for SMD simulations with any of non-contacted subunit pairs, where the maximum decrease is 0.34 in the selection of α_2_β_1_. Then, these accelerating effects on αβ dimer dissociation should be associated with credibly high emergence of ββ dimer under the simulation condition that Δ*d* ranges from 8 to 12 Å.

**Table 5.**
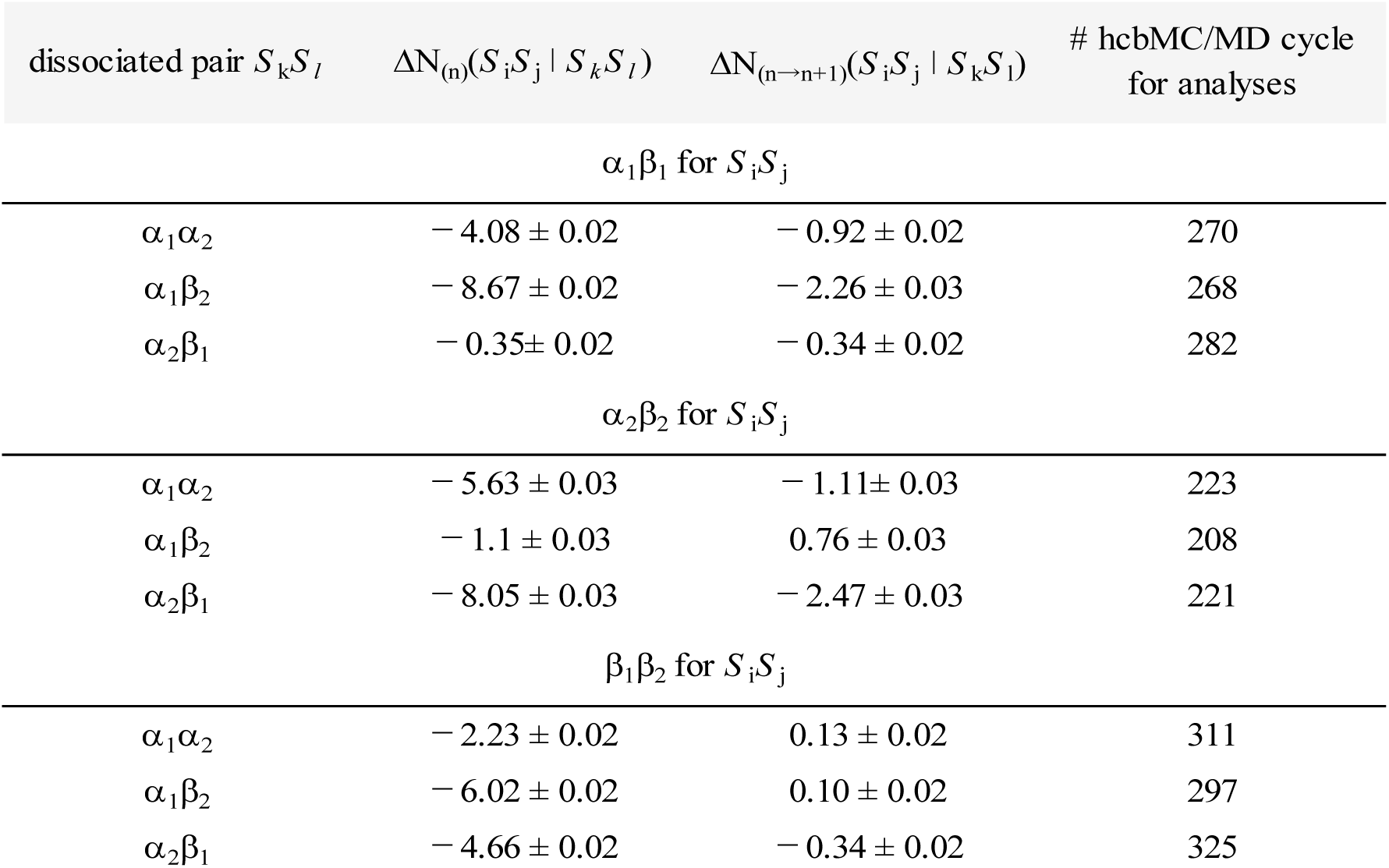
Effects of non-contacted pair dissociation on break and recovery of atomic contact formation between contacted subunit pair. Error estimation denotes 95% confidential interval.

It appears that the RS scheme is similar to the LS one in the performance of subcomplex population prediction. Meanwhile, we will show below the practical advantage of the LS scheme, that is, the robustness of prediction results against selection of an hcbMC/MD simulation condition.

We additionally tested generation of subcomplex populations by using the different setting for increment of inter-COM distance, where Δ*d* is randomly given from the integer ranges between 13 and 17 Å as in the case of the above simulations with the LS scheme. The independent 20 hcbMC/MD simulations were carried out with this setting and resulted in the much lower success rate, 0.55 (see the left column in the second row in **Table 3**, and **Figure S6** for subcomplex generations along the hcbMC/MD cycle). This value of *θ*_ββ_ cannot be statistically distinguished from 0.5 (**Figure 4**). Thus, we can find that the hcbMC/MD simulations with the RS scheme do not necessarily give the MS-consistent disassembly pathway for the tryptophan synthase tetramer.

The simulations with larger Δ*d* speed up dissociations of subunit pairs. This is shown by the fact that the values of χ^dimer^ become smaller in use of larger value range of Δ*d* (see **Table S4**). The accelerated dissociations should be due to larger displacement in each SMD simulations. Such acceleration possibly accompanies greater losses of atomic contacts between a subunit pair and also prevents recovery of the atomic contacts between ββ dimer. Although we employ Metropolis judgement (**3. 1** in **Figure 2A**) with aiming to circumvent unfavorable configurational changes, it does not work to suppress ββ dissociation.

Meanwhile, even if we use the greater value for Δ*d*, which ranges between 18 and 22 Å, the hcbMC/MD simulations with the LS scheme still show high success rates, 0.95 (see the second row in **Table 3**, and **Figure S7** for subcomplex generations along the hcbMC/MD cycle). This clearly shows that the newly proposed LS scheme gives robustness of disassembly process predictions with no regard to an hcbMC/MD simulation condition, increment of inter-COM distance, Δ*d*.

Upon the practical applications of hcbMC/MD approaches, there has been no established protocols to determine the simulation conditions for Δ*d*, force constant for harmonic potential in SMD simulations, and simulation time lengths. In other words, we usually need to give them without *a priori* knowledge for physicochemical properties of multiple protein complex of interest, so far.

As seen above, the RS scheme seems to be substantially affected by a simulation condition such as the range of distance increment. Then, the usage of the LS scheme seems to be useful to stably obtain probable subcomplex populations consistent with MS experiments. We can suppose that this robustness is the practical advantage of the LS scheme upon *ab initio* predictions of disassembly processes of the multimeric protein complex. We could reliably predict such a disassembly process within the context of the hcbMC/MD approach, even if we have no experimental observations for multimeric protein complexes of interest.

### Hybrid MC/MD approach with likelihood-based scheme can predict subcomplex generation of other tetramer complexes

Aiming to test the extensive applicability of our hcbMC/MD approach, we additionally examined other three tetramers: acetyl-CoA carboxylase carboxyltransferase, anthranilate synthase and Co-type nitrile hydratase. The subcomplex populations of these tetramers have been also examined by MS or other biochemical experiments.^6,32,33^

These tetramers are similar to the tryptophan synthase tetramer in the subunit components. Each of them consists of two α subunits and two β subunit. However, the experimentally observed subcomplex is the αβ dimer. Notably, this dimer specie cannot simultaneously appear with the ββ dimer (see **Figures 6**, **7** and **8**). Then, we can evaluate the performance of subcomplex population predictions by considering χ^dimer^ (the hcbMC/MD cycle at which any dimer species first emerges) as in the case of tryptophan synthase tetramer. In all of the hcbMC/MD simulations discussed below, increment of inter-COM distance, Δ*d*, is randomly given from the integer ranges between 18 and 22 Å.

**Figure 6.**
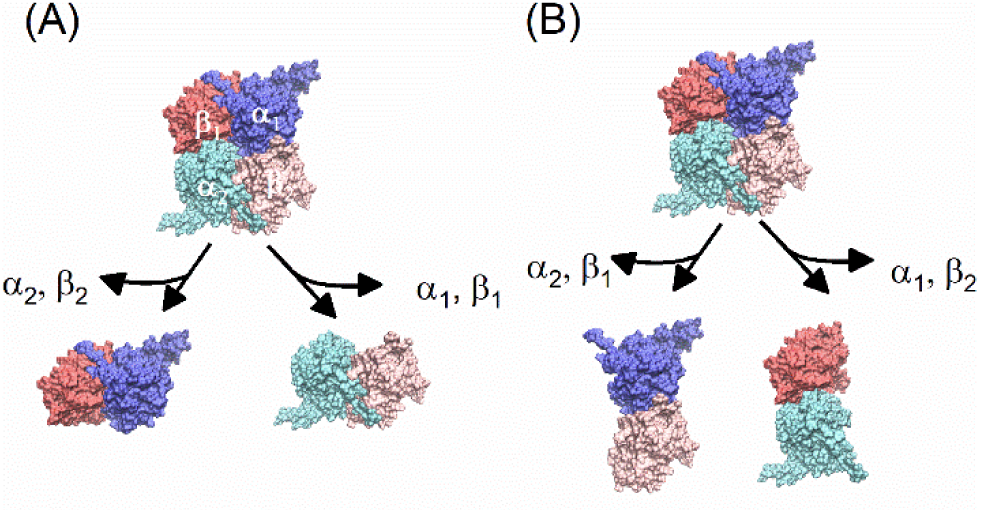
Disassembly pathway of acetyl-CoA carboxylase carboxyltransferase tetramer. (A) α_1_β_1_ or α_2_β_2_ dimer formation. (B) α_1_β_2_ or α_2_β_1_ dimer formation. In the panel A, each of subunits are annotated by greek alphabet with a digit.

**Figure 7.**
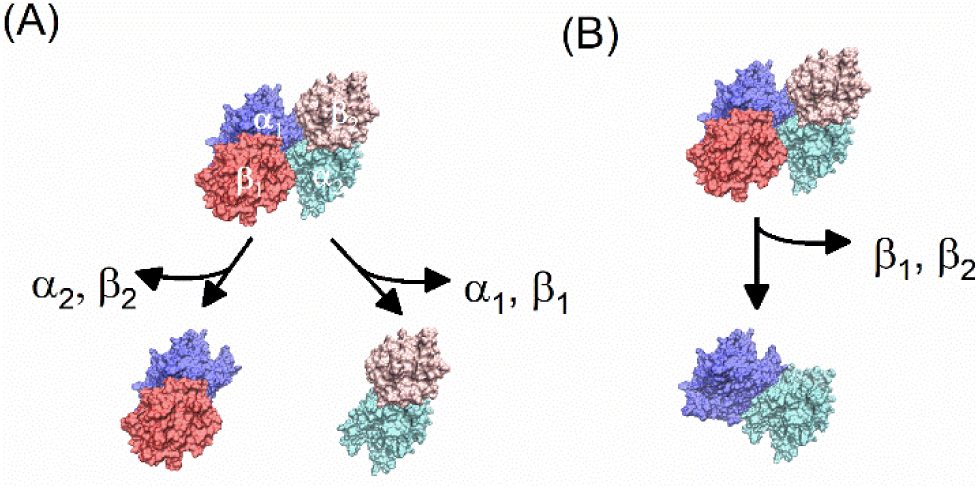
Disassembly pathway of anthranilate synthase tetramer. (A) αβ dimer formation. (B) αα dimer formation. In the panel A, each of subunits are annotated by greek alphabet with a digit.

**Figure 8.**
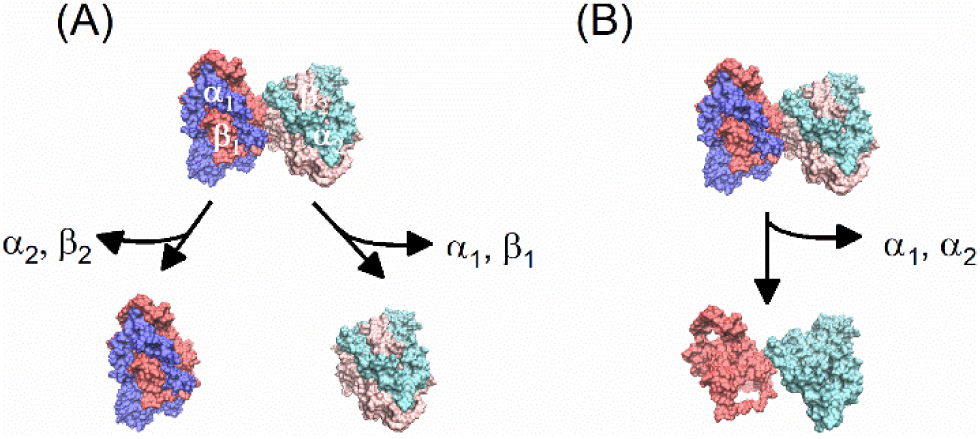
Disassembly pathway of Co-type nitrile hydratase tetramer. (A) αβ dimer formation. (B) ββ dimer formation. In the panel A, each of subunits are annotated by greek alphabet with a digit.

We employed this condition by recalling the robustness of hcbMC/MD simulation results for the tryptophan synthase tetramer. Then, we obtained the high success rate as shown in **Table 6** (The representative configurations are shown in **Figure 3B, 3C** and **3D**; subcomplex generations along the hcbMC/MD cycle are illustrated in **Figures S8, S9** and **S10**). We will remark each of the three systems as follows.

**Table 6.** Success rate of expected dimer emergence for the three tetramers. The number of succeeded and total hcbMC/MD simulations are found at the left and right sides across the slash symbol in the parentheses.

| tetramer | success rate |
| --- | --- |
| Acetyl-coenzyme A carboxylase<br>carboxyltransferase | 1.0 (20/20) |
| Anthranilate synthase | 0.95 (19/20) |
| Co-type nitrile hydratase | 1.0 (20/20) |

The acetyl-CoA carboxylase carboxyltransferase tetramer (ACC) has a circular topology (see **Figure 9B**), while the hcbMC/MD simulations for ACC show high success rate of emergence of the expected dimer (1.0) (see **Table 6**). The α_1_β_1_ and α_2_β_2_ dimers are generated via combination of dissociation between α_1_ and β_2_ and that between α_2_ and β_1_. As shown in **Table 7**, α_1_β_2_ and α_2_β_1_ are selected for dissociation subunit pair with relatively high selection rates, compared with α_1_β_1_. This could simultaneously suppress formation of unexpected dimer, α_1_β_2_ or α_2_β_1_ (**Figure 6**).

**Figure 9.**
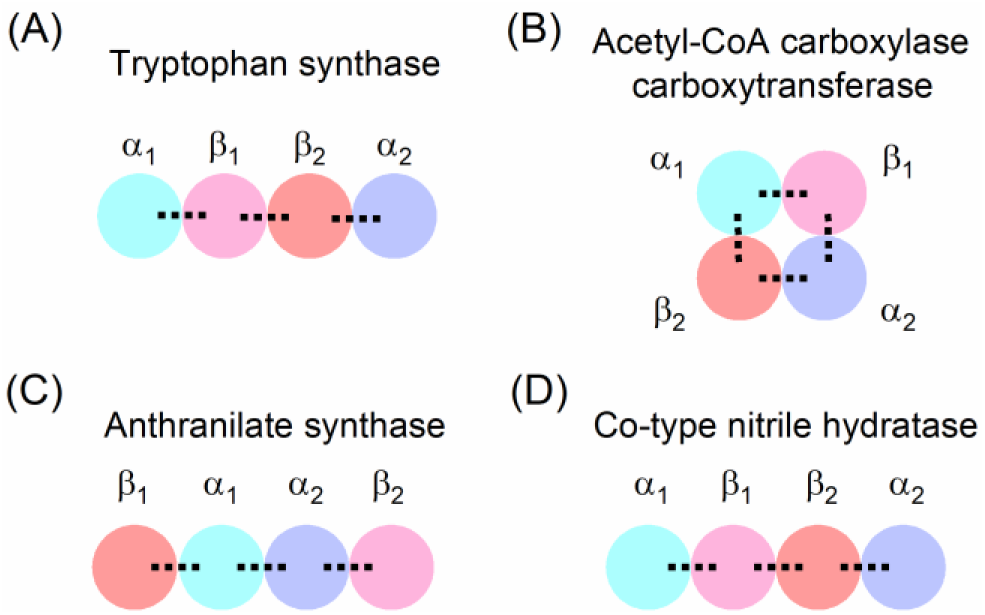
Schematic illustration of inter-subunit interaction topology. (A) Tryptophan synthase. (B) Acetyl-CoA carboxylase carboxytransferase. (C) Anthranilate synthase. (D) Co-type Nitrile hydratase. Dotted lines denote inter-subunit contacts. Salt bridge formations are assigned by considering the initial structures for hybrid configuration bias Monte Carlo/Molecular Dynamics simulations. (see **Table 2**)

**Table 7.** Frequency in percentage for selection of subunit pair for steered MD simulations in hdcMC/MD cycle.

| system | subunit pair |  |  |  |  |  |
| --- | --- | --- | --- | --- | --- | --- |
| | $\alpha_1\beta_1$ | $\alpha_2\beta_2$ | $\alpha_1\alpha_2$ | $\beta_1\beta_2$ | $\alpha_1\beta_2$ | $\alpha_2\beta_1$ |
| Acetyl-coenzyme A carboxylase carboxyl transferase | 9.4 | 26.9 | 7.3 | 0 | 30.3 | 26.1 |
| Anthranilate synthase | 22.6 | 14 | 56.8 | 0 | 2.8 | 3.8 |
| Co-type nitrile hydratase | 8.8 | 36.1 | 0 | 55.1 | 0 | 0 |

Interestingly, 4 of the 20 simulations shows coexistence of αβ dimer and the monomeric subunits (see **Figure S8**). The coexistence of αβ dimer and the two monomers is consistent with the previous MS experiments^5^. The mechanism of monomer emergence could be explained as follows. α_2_β_2_ is similar to α_1_β_2_ and α_2_β_1_ in rates for selection of dissociating subunit pair (see **Table 7**), then leading to release of either α_2_ or β_2_ monomers from the tetramer before generation of the αβ dimer. According to these achievements for the ACC system, we could expect extensive use of our hcbMC/MD approach.

Next, we performed the hcbMC/MD simulations for the anthranilate synthase (AS). AS is also different from the tryptophan synthase in the inter-subunit interaction topology (see **Figure 9A** and **9C**), while the 20 simulations give success rate of 0.95. The AS has two αβ dimers and they make contacts with each of the α subunits. The high frequency of αβ emergence is due to that αα dissociation is preferably selected (see **Table 7**).

This preferable selection could be explained by the structural properties of the initial structure of the AS (see **Table 2**). Among the three subunit pairs, α_1_β_1_, α_2_β_2_, and α_1_α_2_, the number of inter-subunit SB for α_1_α_2_ is the smallest, thus promoting the dissociation of this subunit pair in the simulations. As remarked in Materials and Methods section, we select the initial atomic coordinates for the hybrid MC/MD simulations by considering the average of the number of inter-subunit SB under the thermal fluctuation. From the technical point of view, this step is important to reliably simulate the disassembly in order consistent with MS experimental observations.

It is noted that α_1_β_2_ and α_2_β_1_ are sometime selected as dissociating subunit pair but the initial AS structure does not form salt bridges between α_1_β_2_ and between α_2_β_1_ (see **Table 2**). This is due to transient formation of salt bridge between the subunit pair in hcbMC/MD simulations. Recalling the analyses of salt bridge formation for the unbiased 10 ns-NPT MD simulations for the AS system (see **Table S2**), we could suppose such transient salt bridge formation as a possible microscopic event emerging under thermal fluctuation.

As in the case of the ACC system, the hcbMC/MD simulation for the AS system partly shows the coexistence of the αβ dimer and monomer subunits. The 10 of the 19 successful simulations show such subcomplex populations (see **Figure S9** in Supporting Information). To our best knowledge, there are no previous MS studies for disassembly pathways of the AS (and also for Co-type nitrile hydratase, considered below). Testing this prediction by using MS experiments can give further evaluation of reliability of our hcbMC/MD approach.

Lastly, we examined the Co-type nitrile hydratase (CNH) and observed emergence of the expected dimer species, αβ, for the all of the 20 simulations for CNH (see **Table 6**). This tetramer has the inter-subunit interaction topology similar to tryptophan synthase (See **Figure 9A** and **9D**), while we succeeded to simulate the αβ dimer formation (see **Figure S10** in Supporting Information). The high success rate could be also explained by the structural properties of CNH as in the case of AS. The number of salt bridge formation between β_1_β_2_ are 10-fold smaller than that between α_1_β_1_ and that between α_2_β_2_ in the configuration studied here (see **Table 3**). The significant difference in the number of salt bridge formation contributes to preferable selection of β_1_β_2_ dimer as dissociation pair and then leads to emergence of αβ dimer (see **Table 7**).

In summary, we applied the hcbMC/MD simulations to the four heteromeric tetramers and succeeded to obtain subcomplex populations experimentally observed. Employing likelihood function can lead to the reliable predictions of heteromeic protein complexes with multiple subunits. Meanwhile, recalling the definition given in eqn. 7, the current version of the likelihood function may be specialized to multimeric protein complexes whose energetic stability is finely characterized by the salt-bridge formations. Aiming to further extensively apply our hcbMC/MD approaches to biologically important complexes, such as protein-nucleotide complexes, we will develop a general-purpose likelihood function in future study.

## Concluding Remarks

The physicochemical entity of biological phenomenon in the cell is a network of biochemical reactions and the activity of such a reaction is regulated by a specific set of multimeric protein complexes. Thus, to understand the design principle of the reaction, and furthermore the network of reactions observed in the biological system, it is a critical step to physicochemically characterize formation mechanisms of their assembly and disassembly. However, such observations have not been satisfactorily obtained due to applicable limitations of the pre-existing methodologies, MS experiments and multimeric protein docking simulations based on structural bioinformatics techniques.

To address the problem, we have developed the hcbMC/MD simulations and employed it to examine disassembly processes of four heteromeric protein complexes in this study. By employing the likelihood-based selection scheme, we succeeded in highly reliable predictions of the disassembly orders of these proteins without *a priori* knowledge of MS experiments and structural bioinformatics simulations. Besides, this new simulation procedure reproduced the observations we obtained for the homomeric protein complex. It can be therefore said that the newly proposed procedure is a straightforward improvement of the previous one.

These achievements indicate that our hcbMC/MD simulation approach is available to give a complete set of atomic coordinates for multimeric protein complex systems at any of intermediate states in disassembly processes, that is, a primary information toward further physicochemical analyses of temporal dynamics of the processes (e.g., free energy profile calculations). Due to the complementary roles for the two pre-existing methodologies, our hcbMC/MD simulation can be the third methodology in the proteomics research field.

## Acknowledgements

This work was supported by a Grant-Aid for Scientific Research on Innovative Areas “Chemistry for Multimolecular Crowding Biosystems” (JSPS KAKENHI Grand No. JP17H06353).

