## Supporting Information for "Computational Prediction of Heteromeric Protein Complex Disassembly Order with Hybrid Monte Carlo/Molecular Dynamics Simulation"

---

[a]Department of Computational Science, Graduate School of System Informatics, Kobe University, 1-1 Rokkodai-cho, Nada-ku, Kobe 657-8501, Japan

[*]Ikuo Kurisaki

[*]Shigenori Tanaka

### SI-1. Disassembly of serum amyloid P component protein pentamer (SAP) by using hybrid configuration biased (hcb) MC/MD simulation with likelihood-based selection (LS) scheme.

The molecular components of SAP system are the same as that discussed in our previous study[1]. The initial atomic coordinates for the hcbMC/MD simulations were obtained from the 10 ns-NPT MD simulation as in the case of our previous study. A hcbMC/MD simulation with the LS scheme were independently carried out five times. An increment of inter-COM distance at each hcbMC/MD cycle is sampled from the integer distribution which ranges from 13 to 17 Å as in the case of tryptophan synthase in this study.

**Figure S3** shows the disassembly pathway of SAP pentamer for each of the five simulations. As shown in **Figure S1**, there are two exclusive pathways for initial subunit dissociations. We observed both of the trimer generation pathway (panels A, B, D and E in **Figure S3**) and the tetramer generation pathway (panel C in **Figure S3**).

### SI-2. Analyses of inter-subunit interface in the initial structures for hcbMC/MD simulations

We performed 10 independent 10-ns NPT simulations and calculated RMSd value by referring the X-ray crystallography structure for each system (**Figure S11**) (the technical details for 10-ns NPT simulations are given in **SI-3** and **SI-4**). The RMSd values are averaged at each time point over the 10 simulations. Assuming that the RMSd is converged after 8 ns, we calculated the number of salt-bridge formation between subunits (**Table S2**).

### SI-3. Setup of molecular dynamics simulations

In each simulation, electrostatic interaction was treated by the Particle Mesh Ewald method, where the real space cutoff was set to 9 Å. The vibrational motions associated with hydrogen atoms were frozen by SHAKE algorithm through MD simulations. The translational center-of-mass motion of the whole system was removed by every 500 steps to keep the whole system around the origin, avoiding an overflow of coordinate information from the MD trajectory format. Snapshot structures were recorded by every 10 ps. These simulation conditions referred above were common in all of the simulations discussed in this manuscript.

### SI-4. Preparation of initial atomic coordinates for hcbMC/MD simulations

Each atomic coordinate of water and $K^+$ molecules in a heteromeric protein tetramer system was energetically relaxed by the following molecular mechanics (MM) and molecular dynamics (MD) simulations. First, steric clashes in the system were removed by MM simulation, which consists of 1000 steps of the steepest descent method followed by 49000 steps of the conjugate gradient method.

Then the system temperature and density were relaxed through the following five MD simulations: NVT (0.001 to 1 K, 0.1 ps) → NVT (1 K, 0.1 ps) → NVT (1 to 300 K, 20 ps) → NVT (300 K, 20 ps) → NPT (300 K, 300 ps, 1 bar). In each of MM and MD simulations, the atomic coordinates of non-hydrogen atoms in the tetramer were restrained by the harmonic potential with force constant of 10 kcal/mol/$Å^2$ around the initial atomic coordinates. The first two NVT MD simulations and the other MD simulations were performed using 0.01 fs and 2 fs for the time step of integration, respectively. The first and second NVT MD simulations were performed using Berendsen thermostat[2] with a 0.001 ps of coupling constant. Meanwhile the following three simulations were performed using Langevin thermostat with 1-$ps^{-1}$ of collision coefficient. In the first NVT MD simulation, the reference temperature was linearly increased along the time-course. In the NPT MD simulation, the system pressure was regulated with Monte Carlo barostat, where the system volume change was attempted by every 100 steps. Each set of initial atomic velocities was randomly assigned from the Maxwellian distribution at 0.001 K.

For each atomic coordinates obtained above, a tetramer conformation also was structurally relaxed in aqueous solution through the following 7-step MD simulations: NVT (0.001 to 1 K, 0.1 ps, 10 kcal/mol/Å) → NVT (1 to 300 K, 0.1 ps, 10 kcal/mol/Å) → NVT (300 K, 10 ps, 10 kcal/mol/Å) → NVT (300 K, 40 ps, 5 kcal/mol/Å) → NVT (300 K, 40 ps, 1 kcal/mol/Å) → NVT (300 K, 40 ps) → NPT (300 K, 1 bar, 10 ns). The first two NVT MD simulations and the other MD simulations were performed using 0.01 fs and 2 fs for the time step of integration, respectively. In the first two NVT simulations, the reference temperature was linearly increased along the time-course. In the first 5 steps, non-hydrogen atoms in the tetramer were positionally restrained by the harmonic potential around the initial atomic coordinates. In each NVT simulation, temperature was regulated using Langevin thermostat with 1-ps$^{-1}$ collision coefficient. In the last 10-ns NPT simulation, temperature and pressure were regulated by Berendsen thermostat[2] with a 5-ps coupling constant, and Monte Carlo barostat, where system volume change was attempted by every 100 steps, respectively. The initial atomic velocities were randomly assigned from the Maxwellian distribution at 0.001 K. The snapshot structure obtained from the 10-ns NPT MD simulation procedure was employed for the following hcbMC/MD simulations.

### SI-5. Configuration generation at each hcbMC/MD cycle by using unbiased and steered MD simulations

We here describe the computational details involved with **1.1** and **2.2** in the hcbMC/MD workflow (**Figure 2**). Sets of atomic coordinates of heteromeic protein complex system, denoted by $\{\mathbf{a}_1, \mathbf{a}_2, \cdots \mathbf{a}_M\}$ and $\{\mathbf{b}_1, \mathbf{b}_2, \cdots \mathbf{b}_M\}$ above, were sampled by using an unbiased MD simulation and a steered molecular dynamics (SMD) simulation under the NPT conditions (300 K; 1 bar), respectively.

The simulation length is 100 ps for each of MD and SMD simulations. The system temperature and pressure were regulated by Langevin thermostat with 1-ps$^{-1}$ collision coefficient, and Monte Carlo barostat with attempt of system volume change by every 100 steps, respectively. A set of atomic coordinates was recorded by every 10 ps and was indexed in ascending order to prepare $\{\mathbf{a}_1, \mathbf{a}_2, \cdots \mathbf{a}_M\}$ and $\{\mathbf{b}_1, \mathbf{b}_2, \cdots \mathbf{b}_M\}$; M is set to 10 in the present simulations.

We employed the distance between centers of mass of the chosen subunit pair as the reaction coordinate of SMD simulation. A COM for a subunit was calculated by using the all of $C_\alpha$ atoms. For such relatively large subunits which including more than 256 $C_\alpha$ atoms, we used the half of the $C_\alpha$ atoms, which are assigned by odd numbers in each subunit.

In an SMD simulation, a target value of the distance was set to $d_0 + \Delta d$, where $d_0$ and $\Delta d$ are an initial value of the distance calculated from the $\mathbf{x}_o$ at the cycle and a random integer, respectively. We used two different integer range to give $\Delta d$: one is the range of 8 to 12, and the other is the range of 13 to 17. The harmonic potential with the force constant of 10 kcal/mol/Å$^2$ was imposed on the reaction coordinate as in the case of our previous study. In each cycle, random seeds were independently given. Additional computational setups are similar to those described in **SI-3** in Supporting Information, respectively.

### SI-6. Mechanisms of progression of ββ dimer dissociation

Considering the two hcbMC/MD simulations which failed to generate ββ dimer, we can find how dissociation of ββ dimer progresses. Firstly, the number of salt bridge formation between the β subunits is transiently decrease, then being close to, or smaller than that between α and β subunits. As for the panels A and B in **Figure S12**, 50th and 63th cycles are assigned to such conditions, respectively.

We can estimate the effect of difference of $N_{SB}$ on likelihood of selecting subunit pair, by considering αββ trimer (see **Figure 1**C). Representative situations are summarized in **Table S5**. As shown in **Table S5**, $N_{SB}$ of ββ is just greater by 1 than that of αβ and probability of selecting ββ is 16% (the 50th cycle in **Figure S12A** is an example), then being probable to be selected in the probabilistic process. In contrast, if $N_{SB}$ of ββ is just smaller by 1 than that of αβ dimer, probability of selecting ββ dimer is 84% (the 39th cycle in **Figure S12A** is an example), then being probable to be selected in the probabilistic process.

Once a ββ dissociating configuration is accepted for the next cycle via Metropolis algorism at each of these cycles, ββ dimer still would have higher likelihood for dissociation subunit pair. Then, the break of salt bridge between ββ dimer enhance to dissociation of ββ dimer. We can find consecutive decrease of atomic contact between ββ dimer (**Figure S12C** and **S12D**).

As remarked in the main text, the above situations could happen via thermal fluctuation, which is originated from intrinsic randomness of assigning initial atomic velocities at each simulation cycle and temperature regulation with a probabilistic thermostat. Nonetheless, the ration of failed simulations with the LS scheme is 0.1 (only two cases in the twenty simulations) so that we suppose that the effects of thermal fluctuation is satisfactorily suppressed in the present simulations and does not have significant influence on prediction of subcomplex populations.

**Table S1**. Amino acid sequences of subunits consisting of the tetramers.

| system (abbreviated name) | amino acid sequence | |
|---|---|---|
| | α subunit | β subunit |
| Tryptophan Synthase (TS) | MERYENLFAQLNDRREGAFVPFVTLGDP<br>GIEQSLKIIDTLIDAGADALELGVPFSD<br>PLADGPTIQNANLRAFAAGVTPAQCFEM<br>LALIREKHPTIPIGLLMYANLVFNNGID<br>AFYARCEQVGVDSVLVADVPVEESAPFR<br>QAALRHNIAPIFICPPNADDDLLRQVAS<br>YGRGYTYLLSRSGVTGAENRGALPLHHL<br>IEKLKEYHAAPALQGFGISSPEQVSAAV<br>RAGAAGAISGSAIVKIIEKNLASPKQML<br>AELRSFVSAMKAASRA | TTLLNPYFGEFGGMYVPQILMPALNQLE<br>EAFVSAQKDPEFQAQFADLLKNYAGRPT<br>ALTKCQNITAGTRTTLYLKREDLLHGGA<br>HKTNQVLGQALLAKRMGKSEIIAETGAG<br>QHGVASALASALLGLKCRIYMGAKDVER<br>QSPNVFRMRLMGAEVIPVHSGSATLKDA<br>CNEALRDWSGSYETAHYMLGTAAGPHPY<br>PTIVREFQRMIGEETKAQILDKEGRLPD<br>AVIACVGGGSNAIGMFADFINDTSVGLI<br>GVEPGGHGIETGEHGAPLKHGRVGIYFG<br>MKAPMMQTADGQIEESYSISAGLDFPSV<br>GPQHAYLNSIGRADYVSITDDEALEAFK<br>TLCRHEGIIPALESSHALAHALKMMREQ<br>PEKEQLLVVNLSGRGDKDIFTVHDILKA<br>RG |
| Acetyl-coenzyme A carboxylase carboxyl transferase (ACT) | MLDFEKPLFEIRNKIESLKESQDKNDVD<br>LQEEIDMLEASLERETKKIYTNLKPWDR<br>VQIARLQERPTTLDYIPYIFDSFMELHG<br>DRNFRDDPAMIGGIGFLNGRAVTVIGQQ<br>RGKDTKDNIYRNFGMAHPEGYRKALRLM<br>KQAEKFNRPIFTFIDTKGAYPGKAAEER<br>GQSESIATNLIEMASLKVPVIAIVIGEG<br>GSGGALGIGIANKVLMLENSTYSVISPE<br>GAAALLWKDSNLAKIAAETMKITAHDIK<br>QLGIIDDVISEPLGGAHKDIEQQALAIK<br>SAFVAQLDSLESLSRDEIANDRFEKFRN<br>IGSYIE | IMTKCPKCKKIMYTKELAENLNVCFNCD<br>HHIALTAYKRIEAISDEGSFTEFDKGMT<br>SANPLDFPSYLEKIEKDQQKTGLKEAVV<br>TGTAQLDGMKFGVAVMDSRFRMGSMGSV<br>IGEKICRIIDYCTENRLPFILFSASGGA<br>RMQEGIISLMQMGKTSVSLKRHSDAGLL<br>YISYLTHPTTGGVSASFASVGDINLSEP<br>KALIGFAGRRVIEQTINEKLPDDFQTAE<br>FLLEHGQLDKVVHRNDMRQTLSEILKIH<br>QEV |
| Co-type nitrile hydratase (CNH) | DHHHDGYQAPPEDIALRVKALESLLIEK<br>GLVDPAAMDLVVQTYEHKVGPRNGAKVV<br>AKAWVDPAYKARLLADGTAGIAELGFSG<br>VQGEDMVILENTPAVHNVFVCTLCSCYP<br>WPTLGLPPAWYKAAPYRSRMVSDPRGVL<br>AEFGLVIPANKEIRVWDTTAELRYMVLP<br>ERPAGTEAYSEEQLAELVTRDSMIGTGL<br>PTQP | MNGIHDTGGAHGYGPVYREPNEPVFRYD<br>WEKTVMSLLPALLANGNFNLDEFRHSIE<br>RMGPAHYLEGTYYEHWLHVFENLLVEKG<br>VLTATEVATGKAASGKTATPVLTPAIVD<br>GLLSTGASAAREEGARARFAVGDKVRVL<br>NKNPVGHTRMPRYTRGKVGTVVIDHGVF<br>VTPDTAAHGKGEHPQHVYTVSFTSVELW<br>GQDASSPKDTIRVDLWDDYLEPA |
| Anthranilate synthase (AS) | TKPQLTLLKVQASYRGDPTTLFHQLCGA<br>RPATLLLESAEINDKQNLQSLLVIDSAL<br>RITALGHTVSVQALTANGPALLPLLDEA<br>LPPEVRNQARPNGRELTFPAIDAVQDED<br>ARLRSLSVFDALRTILTLVDSPADEREA<br>VMLGGLFAYDLVAGFENLPALRQDQRCP<br>DFCFYLAETLLVLDHQRGSARLQASVFS<br>EQASEAQRLQHRLEQLQAELQQPPQPIP<br>HQKLENMQLSCNQSDEEYGAVVSELQEA<br>IRQGEIFQVVPSRRFSLPCPAPLGPYQT<br>LKDNNPSPYMFFMQDDDFTLFGASPESA<br>LKYDAGNRQIEIYPIAGTRPRGRRADGS<br>LDLDLDSRIELEMRTDHKELAEHLMLVD<br>LARNDLARICQAGSRYVADLTKVDRYSF<br>VMHLVSRVVGTLRADLDVLHAYQACMNM<br>GTLSGAPKVRAMQLIAALRSTRRGSYGG<br>RVGYFTAVRNLDTCIVIRSAYVEDGHRT<br>VQAGAGVVQDSIPEREADETRNKARAVL<br>RAIATAHHAKEVF | ADILLLDNVDSFTYNLVDQLRASGHQVV<br>IYRNQIGAEVIIERLQHMEQPVLMLSPG<br>PGTPSEAGCMPELLQRLRGQLPIIGICL<br>GHQAIVEAYGGQVGQAGEILHGKASAIA<br>HDGEGMFAGMANPLPVARYHSLVGSNIP<br>ADLTVNARFGEMVMAVRDDRRRVCGFQF<br>HPESILTTHGARLLEQTLAWALAK |

**Table S2**. Inter-subunit salt bridge formation for the four tetramers. These average values are calculated by using time domains between 8 to 10 ns, which are obtained from 10 independent 10-ns NPT simulations (see **Figure S11** for convergence of RMSd). A stander error was calculated over the 10 independent simulations and the error estimation denotes 95% confidential interval.

| tetramer | subunit pair | | | | | |
|---|---|---|---|---|---|---|
| | $\alpha_1\beta_1$ | $\alpha_2\beta_2$ | $\alpha_1\alpha_2$ | $\beta_1\beta_2$ | $\alpha_1\beta_2$ | $\alpha_2\beta_1$ |
| Tryptophan Synthase | 4.8 ± 0.5 | 4.6 ± 0.4 | 0 | 10.8 ± 2.2 | 0 | 0 |
| Co-type nitrile hydratase | 30.9 ± 2.5 | 0 | 0 | 0 | 2.0 ± 1.3 | 27.7 ± 2.8 |
| Acetyl-coenzyme A carboxylase carboxyl transferase | 14.7 ± 1.1 | 13.7 ± 1.7 | 0 | 0 | 13.0 ± 0.7 | 13.1 ± 0.5 |
| Anthranilate synthase | 17.9 ± 3.2 | 17.2 ± 2.5 | 15.4 ± 1.4 | 0 | 0.1 ± 0.2 | 0.1 ± 0.2 |

**Table S3**. Selection frequency of subunit pair for SMD simulations with the random selection scheme. The hcbMC/MD simulation data are picked up by the first dimer emergence cycle and employed for the analyses. Frequency is shown in percentage.

| distance increase along the reaction coordinate | $\alpha_1\beta_1$ | $\alpha_2\beta_2$ | $\beta_1\beta_2$ | $\alpha_1\alpha_2$ | $\alpha_1\beta_2$ | $\alpha_2\beta_1$ |
|---|---|---|---|---|---|---|
| 8 to 12 Å | 16.7 | 16.9 | 16.0 | 16.6 | 16.6 | 17.2 |
| 13 to 17 Å | 15.2 | 18.2 | 15.3 | 17.2 | 16.6 | 17.5 |

**Table S4**. The cycle when ββ dimer first emerges in simulations for tryptophan synthase tetramer system. The average values are calculated by using all hcbMC/MD simulation trajectories leading to a dimer emergence. Error bars denote 95% confidential interval.

| distance increase along the reaction coordinate | random selection scheme | likelihood-based scheme |
|---|---|---|
| 8 to 12 Å | 235.7 ± 42.4 | - |
| 13 to 17 Å | 52.6 ± 12.6 | 106.8 ± 48.6 |
| 18 to 23 Å | - | 45.7 ± 20.1 |

**Table S5**. Effects of difference of salt bridge formation ($N_{SB}$) between two subunit pair, x and y, on the likelihood of selection.

| $\Delta N_{SB} = N_{SB}(x) - N_{SB}(y)$ | likelihood of selection in % | |
|---|---|---|
| | x | y |
| 0 | 50 | 50 |
| 1 | 15.9 | 84.1 |
| 2 | 3.4 | 96.6 |
| 3 | 0.7 | 99.3 |
| 4 | 0.1 | 99.9 |
| 5 | 0 | 100 |

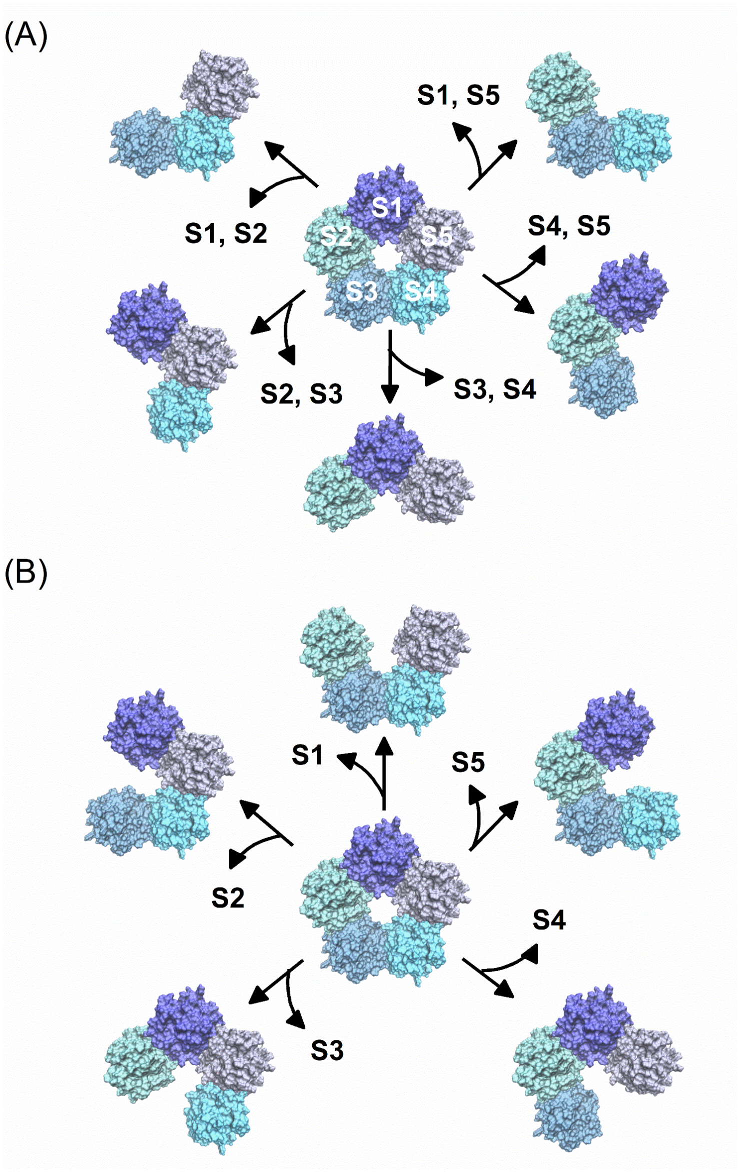


**Figure S1**. The earlier stage of disassembly of serum amyloid P component protein (SAP) pentamer. (A) Trimer formation via dimer dissociation. (B) Tetramer formation via monomer dissociation. Each of SAP subunits are annotated by greek alphabet with a digit.

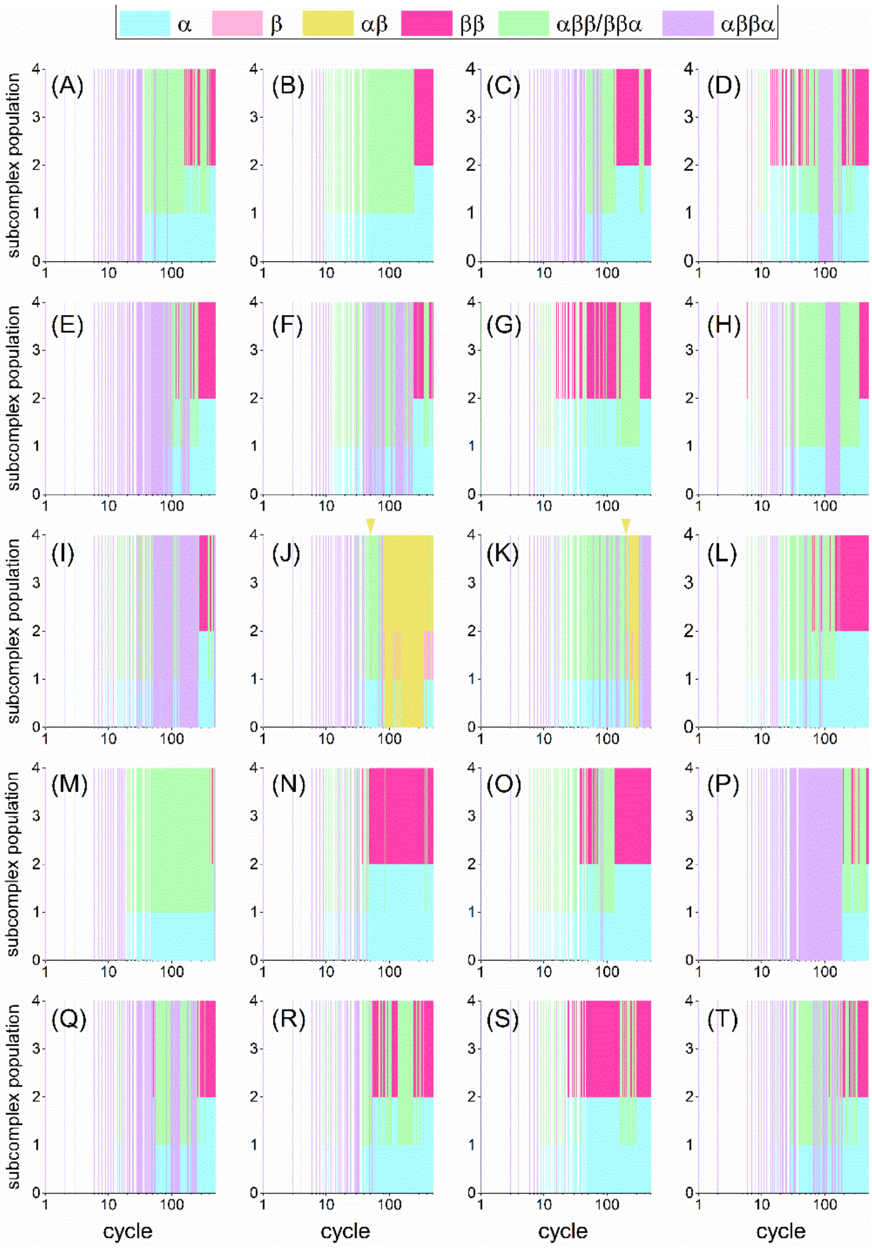


**Figure S2**. 20 independent hcbMC/MD simulations of tryptophan synthase tetramer with the likelihood-based selection scheme by using increment of inter-COM distance for SMD simulations (Δ*d*) which ranges from 13 to 17 Å. Each simulation is annotated by alphabetical character, ranging from A to T. The inverse triangle colored by yellow denotes the cycle when an αβ dimer first emerges during the simulation.

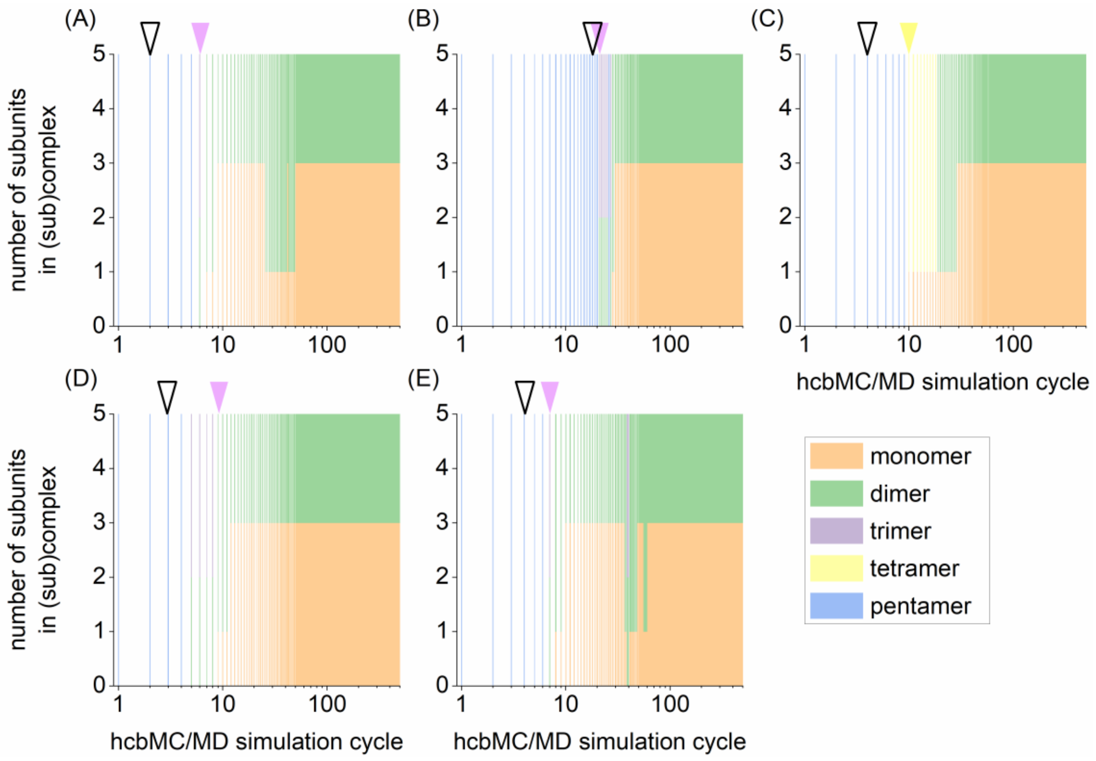


**Figure S3**. 5 independent hcbMC/MD simulations of serum amyloid P component (SAP) homo-pentamer with the likelihood-based selection scheme. The five individual simulations are annotated by A, B, C, D and E. A distribution of SAP (sub)complex species is illustrated by color. Colored and open arrows on panel indicate the initial stage of SAP disassembly (generation of tetramer or trimer) and the first cycle where SAP pentamer undergoes a ring-opening event, respectively. The color of arrow corresponds to the type of larger subcomplex species. Increment of inter-COM distance for SMD simulations ($\Delta d$) ranges from 13 to 17 Å.

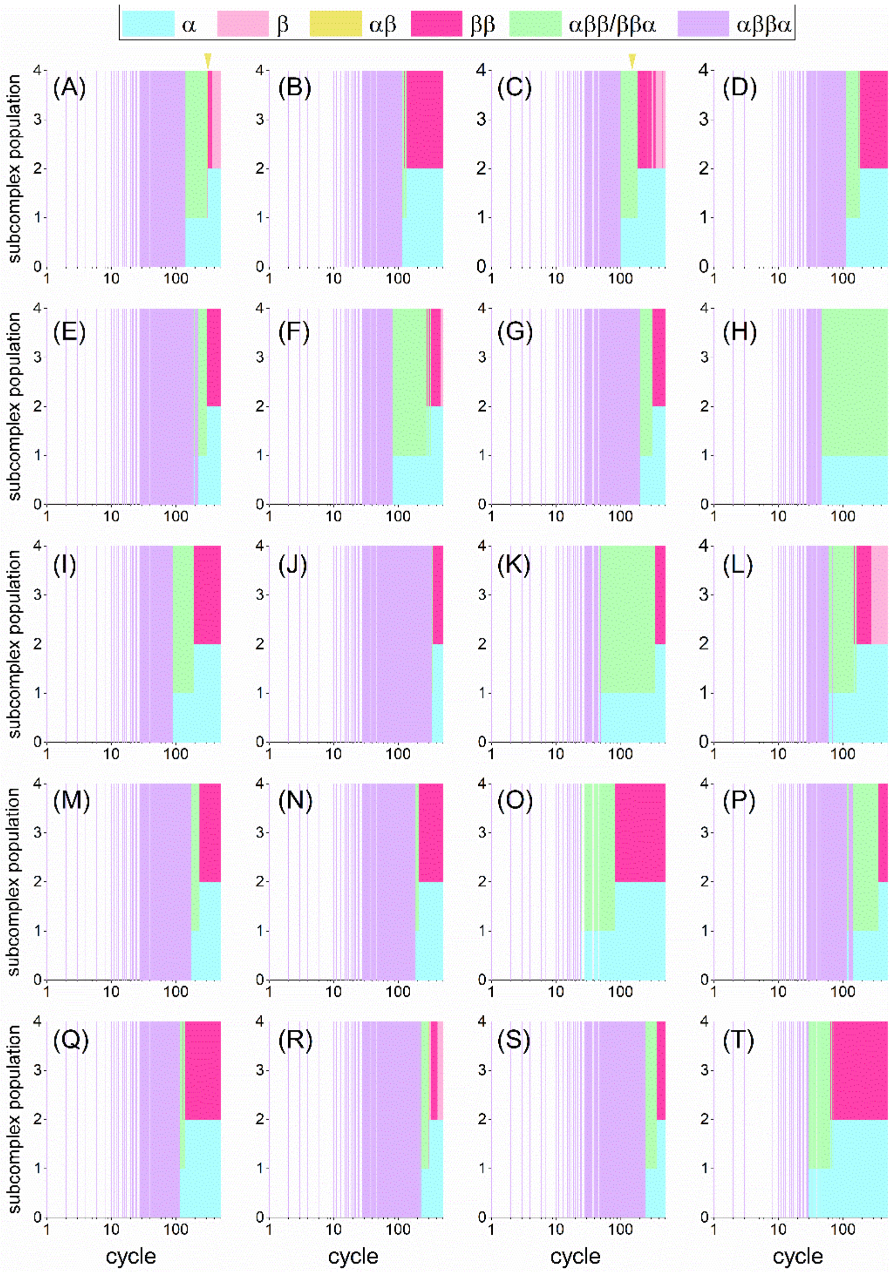


**Figure S4**. 20 independent hcbMC/MD simulations of tryptophan synthase tetramer with the random selection scheme by using increment of inter-COM distance for SMD simulations (Δ*d*) which ranges from 8 to 12 Å. Each simulation is annotated by alphabetical character, ranging from A to T. The inverse triangle colored by yellow denotes the cycle when an αβ dimer first emerges during the simulation.

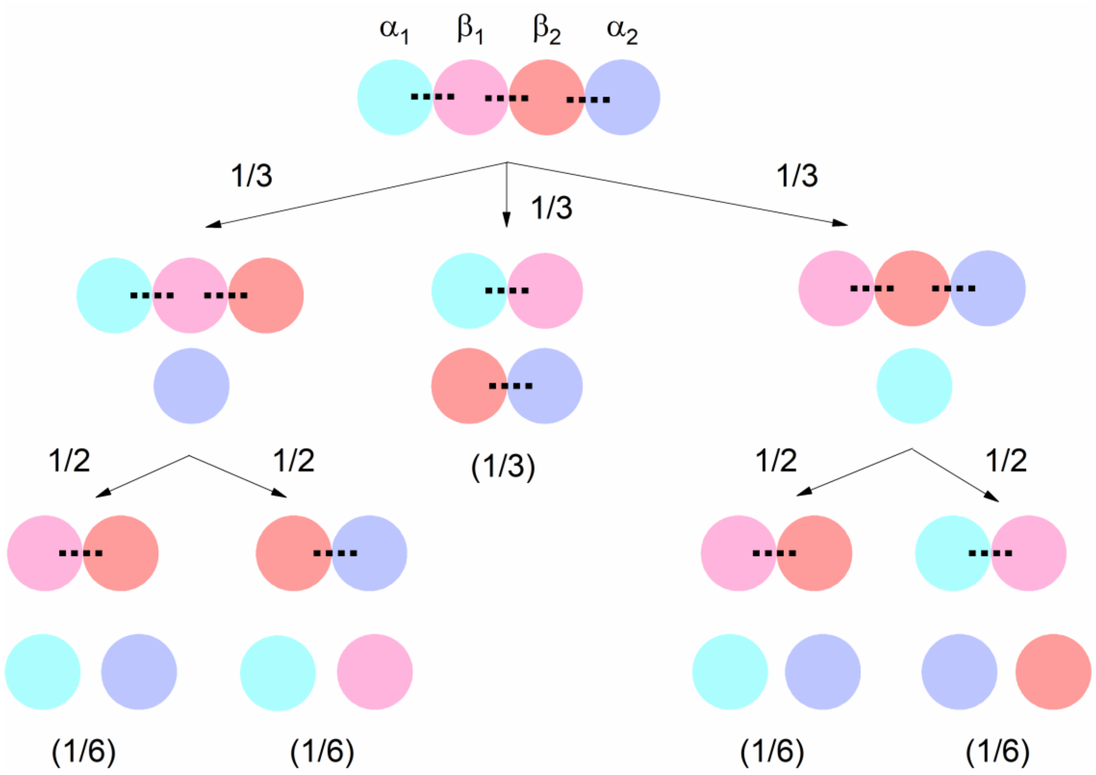


**Figure S5**. Generation probability of subcomplex population of tryptophan synthase. Each selection probability of dissociation pair is assumed to be equal, corresponding to a fraction besides an arrow head. A fraction inside parentheses denotes probability of subcomplex generation.

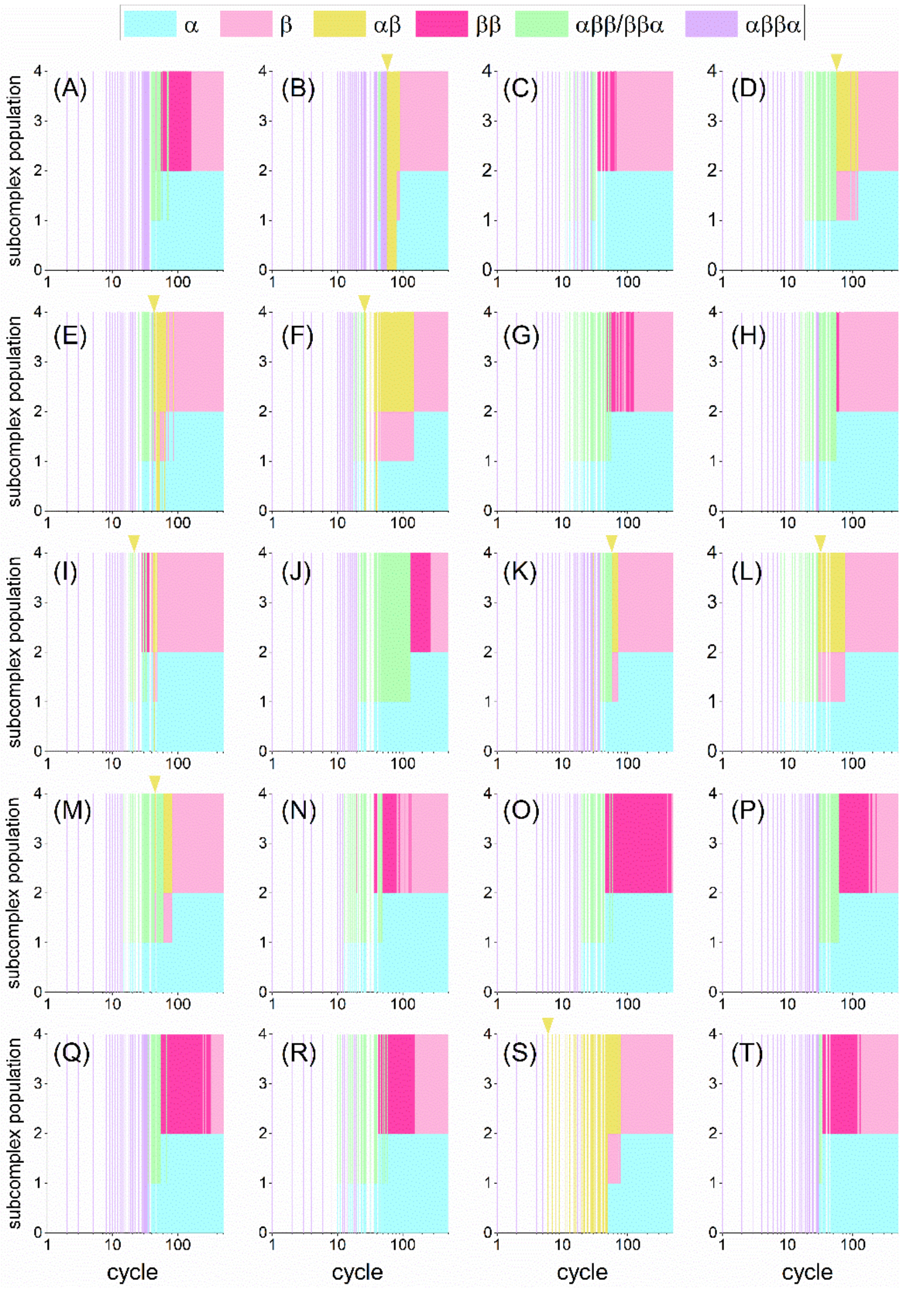


**Figure S6**. 20 independent hcbMC/MD simulations of tryptophan synthase tetramer with the random selection scheme by using increment of inter-COM distance for SMD simulations ($\Delta d$) which ranges from 13 to 17 Å. Each simulation is annotated by alphabetical character, ranging from A to T. The inverse triangle colored by yellow denotes the cycle when an αβ dimer first emerges during the simulation.

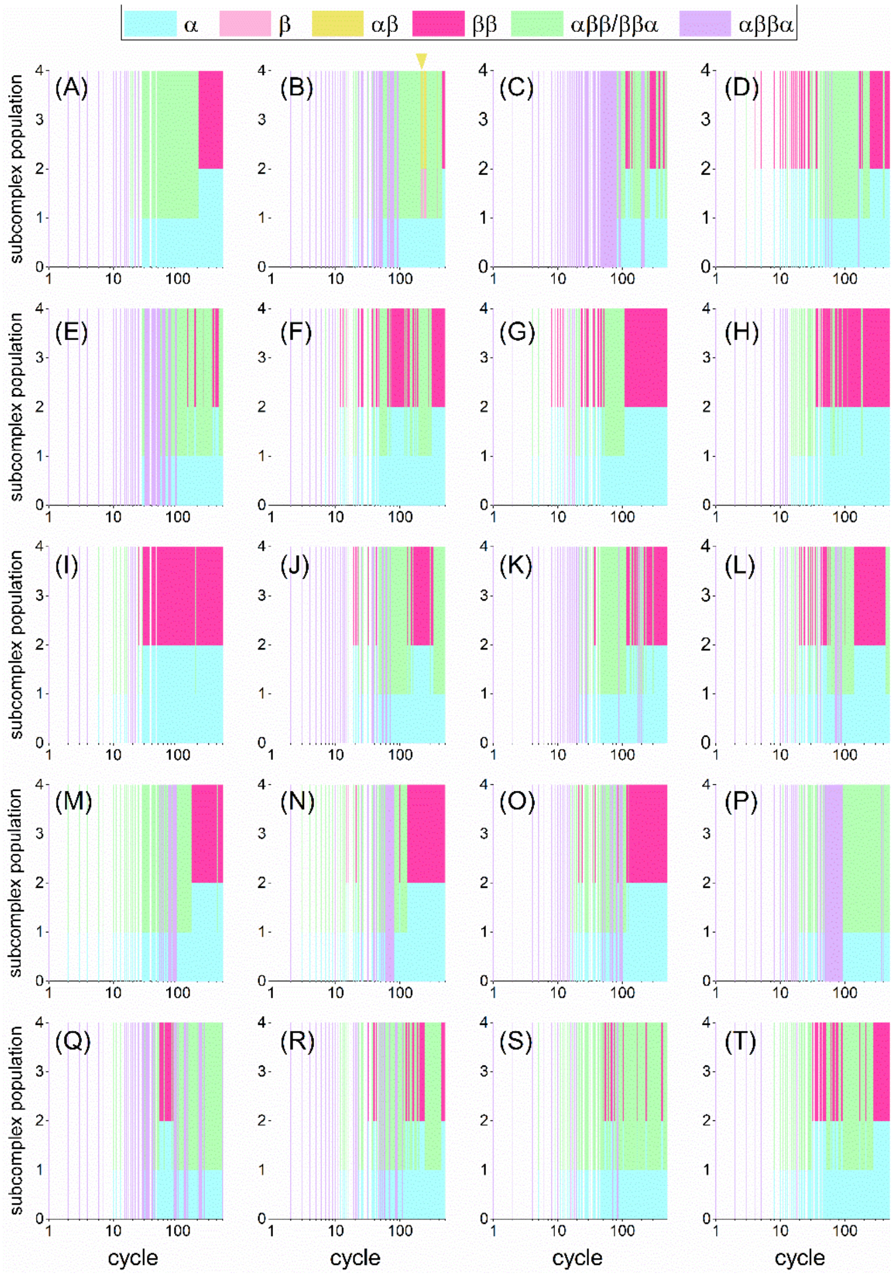


**Figure S7**. 20 independent hcbMC/MD simulations of tryptophan synthase tetramer with the likelihood-based selection scheme by using increment of inter-COM distance for SMD simulations (Δ*d*) which ranges from 18 to 22 Å. Each simulation is annotated by alphabetical character, ranging from A to T. The inverse triangle colored by yellow denotes the cycle when an αβ dimer first emerges during the simulation.

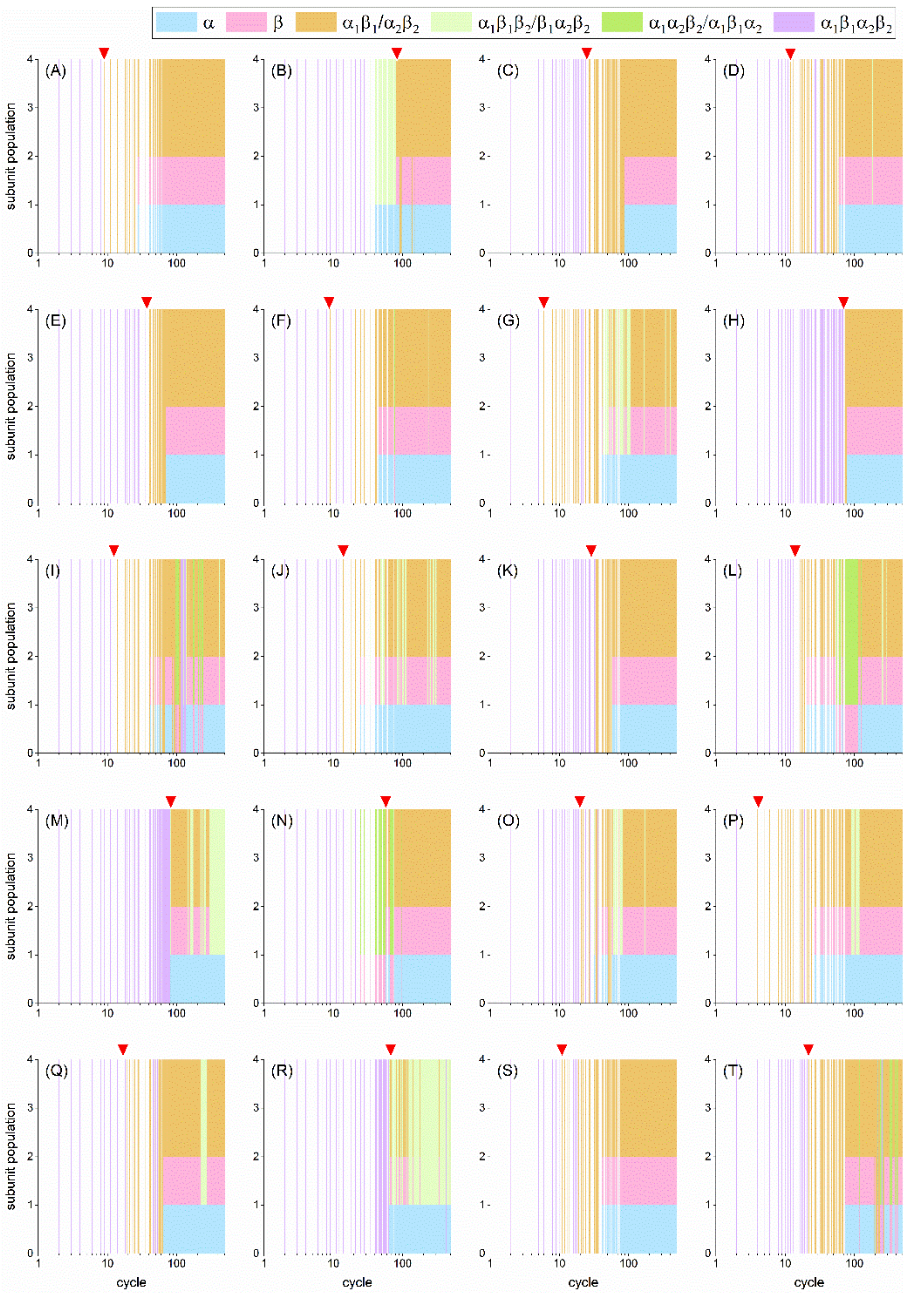


**Figure S8**. 20 independent hcbMC/MD simulations of acetyl-CoA carboxylase carboxyltransferase tetramer with the likelihood-based selection scheme by using increment of inter-COM distance for SMD simulations (Δ*d*) which ranges from 18 to 22 Å. A red arrow on panel indicate the timing of first emergence of dimeric subcomplex. Each simulation is annotated by alphabetical character, ranging from A to T.

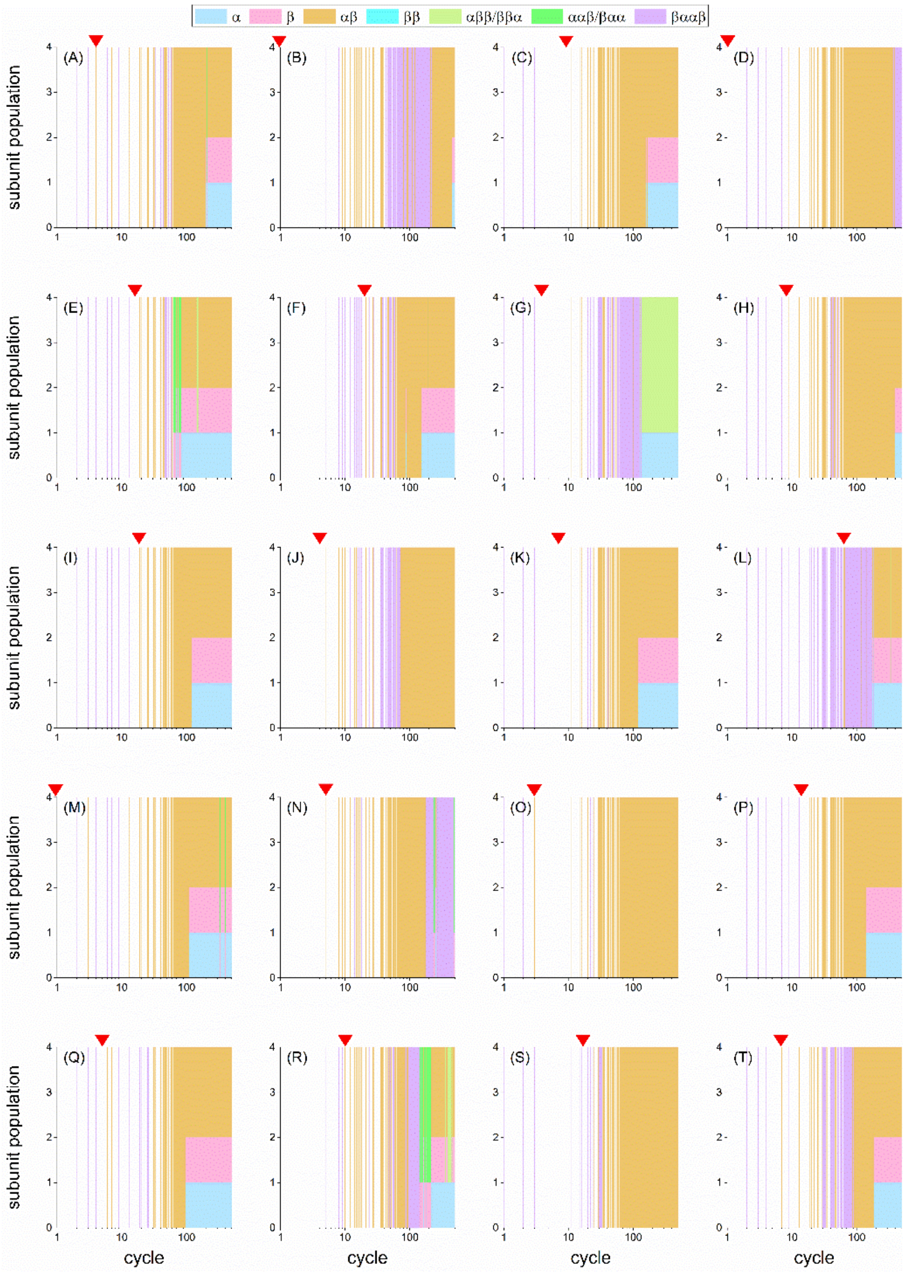


**Figure S9**. 20 independent hcbMC/MD simulations of anthranilate synthase tetramer with the likelihood-based selection scheme by using increment of inter-COM distance for SMD simulations (Δ*d*) which ranges from 18 to 22 Å. A red arrow on panel indicate the timing of first emergence of dimeric subcomplex. Each simulation is annotated by alphabetical character, ranging from A to T.

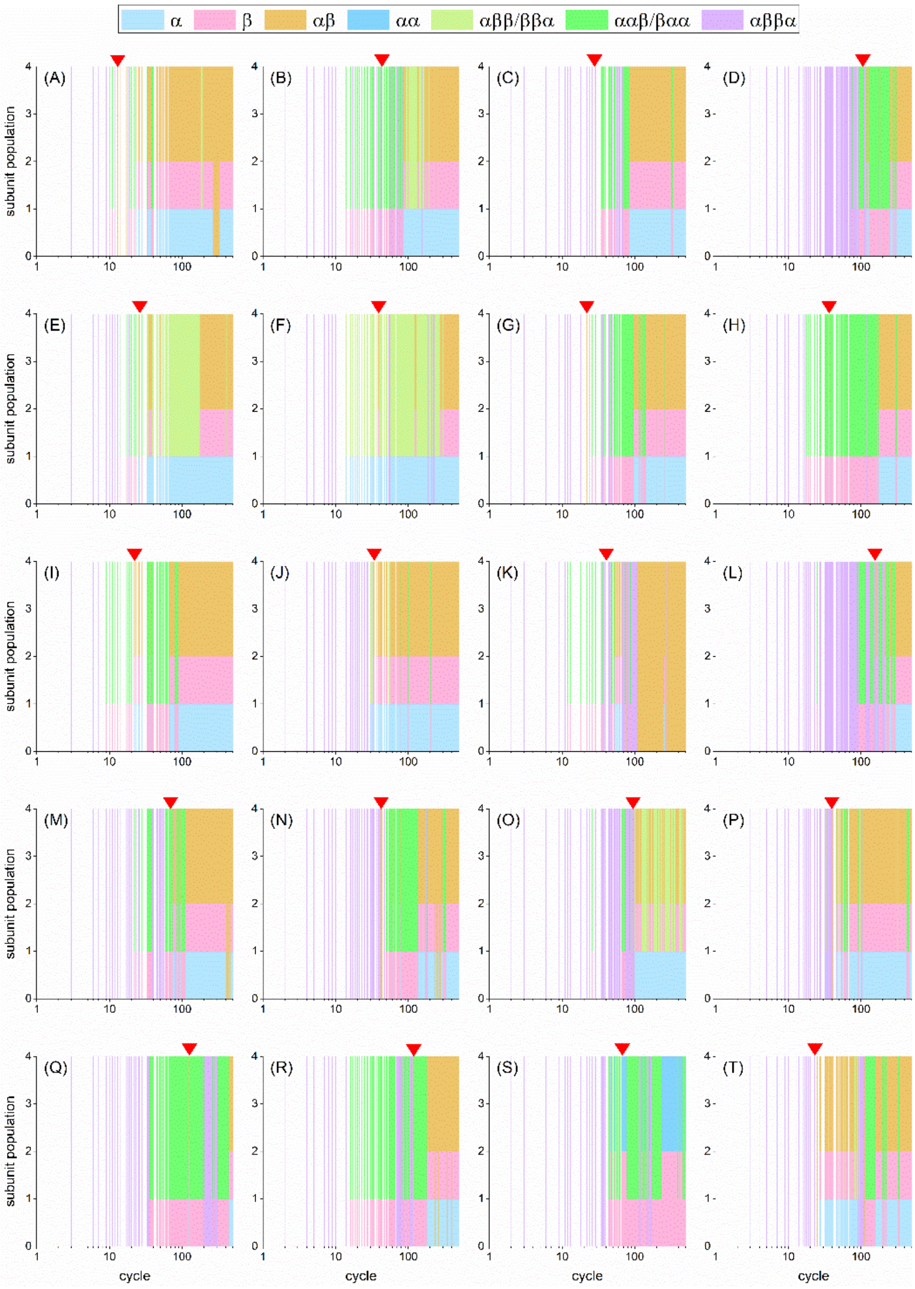


**Figure S10**. 20 independent hcbMC/MD simulations of Co-type nitrile hydratase tetramer with the likelihood-based selection scheme by using increment of inter-COM distance for SMD simulations (Δ*d*) which ranges from 18 to 22 Å. A red arrow on panel indicate the timing of first emergence of dimeric subcomplex. Each simulation is annotated by alphabetical character, ranging from A to T.

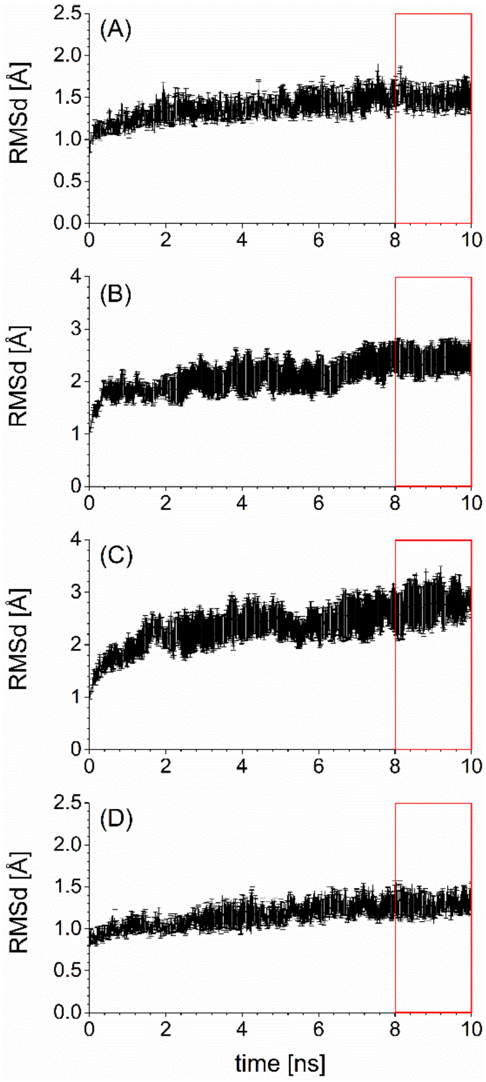


**Figure S11.** Time course change of averaged RMSd for each of the four tetramers. (A) Tryptophan synthase. (B) Acetyl-CoA carboxylase carboxytransferase. (C) Anthranilate synthase. (D) Co Nitrile hydratase. The time domains assumed as convergence are indicated by the red rectangles. These values are calculated by using time domains between 8 to 10 ns, which are obtained from 10 independent 10-ns NPT simulations. An error was calculated over the 10 independent simulations and denotes 95% confidential interval.

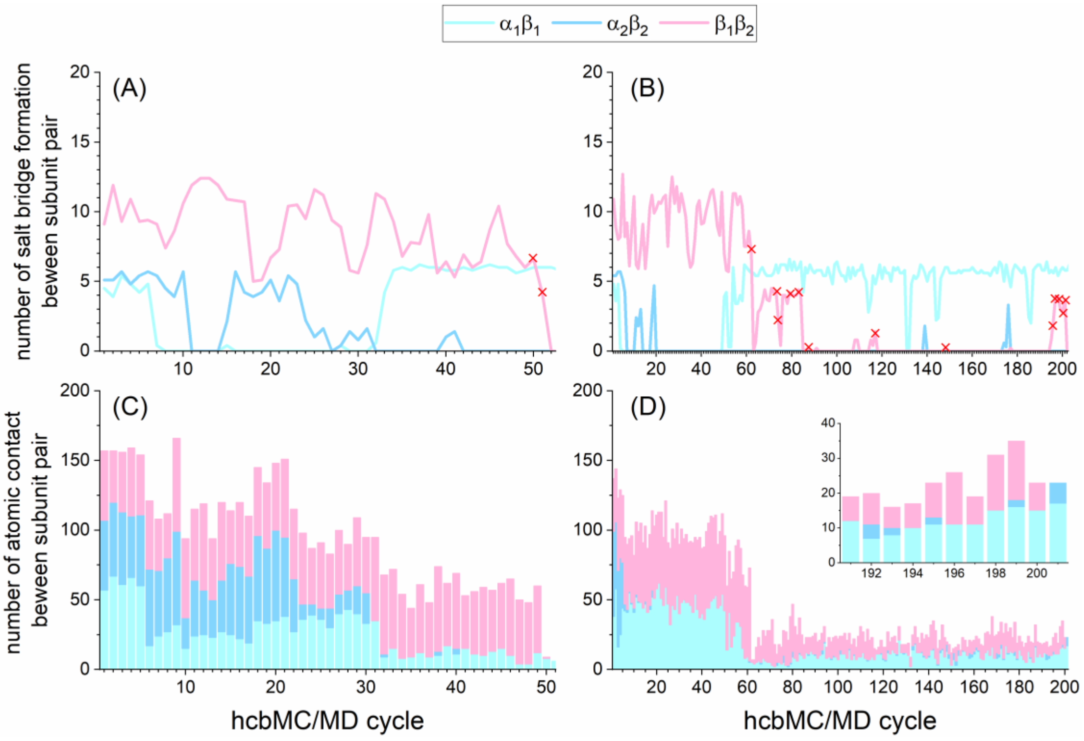


**Figure S12**. Temporal change of salt bridge (SB) formation and atomic contact between subunit pairs of tryptophan synthase for the two simulations which failed to generate MS-observed dimer (annotated by $F_1$ and $F_2$, hereafter). (A) and (B), the number of SB formation for $F_1$ and $F_2$, respectively. The SB number is calculated in average over an MD simulation at the cycle. (C) and (D), the number of atomic contact for $F_1$ and $F_2$, respectively. The value is calculated by using a snapshot structure obtained from the cycle. In the panels A and B, a red cross denotes a simulation cycle where $\beta_1\beta_2$ dissociating SMD trajectory is accepted via Metropolis algorism. In the panel D, the inset highlighted the 190$^{th}$ cycle or later.
